# Time-resolved Interaction Proteomics of the Putative Scaffold Protein GIGANTEA in *Arabidopsis thaliana*

**DOI:** 10.1101/162271

**Authors:** Johanna Krahmer, Greg Goralogia, Akane Kubota, Richard S. Johnson, Young Hun Song, Michael J. MacCoss, Thierry LeBihan, Karen J Halliday, Takato Imaizumi, Andrew J. Millar

## Abstract

The large, plant-specific protein GIGANTEA (GI) is involved in many physiological processes, mediating rhythmic, post-translational regulation in part through circadian and light regulation of *GI* RNA expression. GI binds several proteins implicated in the circadian clock, the control of photoperiodic flowering, and abiotic stress responses, and has co-chaperone activity. By extension, further interaction partners might mediate the less well-understood roles of GI but the number and rhythmicity of these interactors is unknown. Here, we seek potential interactors in a time-specific manner, using quantitative proteomics from a time series study of transgenic *Arabidopsis thaliana* plants constitutively expressing an epitope-tagged GI protein. Previously-identified, direct and indirect interactors of GI were detected but no further F-box proteins related to known GI partners ZTL/FKF1/LKP2. The predominantly non-rhythmic, interacting proteins were implicated in protein folding or degradation, metabolism and chromatin modification, including a small set of partners shared with other clock-related proteins. A transcription factor homologue that we name *CYCLING DOF FACTOR 6* (*CDF6*) was shown to interact both with GI and the ZTL/FKF1/LKP2 proteins and to control photoperiodic flowering. Our results indicate the biochemical pathways, beyond circadian and flowering regulation, that might be affected by GIGANTEA’s rhythmic, post-translational control.

**Significance Statement:** Significance statement of up to two sentences of no more than 75 words total;

The GIGANTEA protein of Arabidopsis was known for circadian and flowering functions, mediated by the FKF1/LKP2/ZTL family of GI-interacting, F-box proteins, then for a co-chaperone activity of unknown scope. We performed time-resolved, interaction proteomics, identifying CDF6 (At1g26790) as a morning-specific GI interactor that controls flowering time. Unlike FKF1 and CDF proteins, most of the 240 candidate partners were not rhythmically enriched. They link GI to proteostasis and metabolic functions that might mediate GI’s physiological functions.

## Introduction

*Arabidopsis thaliana* has well-documented, 24-hour rhythms in many physiological processes, from the elongation rate of seedling hypocotyls to photosynthetic functions and the defense of mature leaves against herbivore attack (Dodd *et al.* 2014, Millar 2016). The overt, circadian rhythms are thought to be driven by a transcriptional-translational feedback circuit (Millar 2016). Detailed, dynamic models based mostly upon transcriptional repression recapitulate the rhythmic expression profiles of these so-called clock genes, including manipulations of the system in mutant plants and under changing photoperiods (Flis *et al.* 2015, Flis *et al.* 2016, Pokhilko *et al.* 2012).

Gene expression switches can operate on a timescale of minutes, however, which does not obviously explain the slow, 24-hour timescale of the circadian clock. Chromatin modification might extend the timescale of transcriptional regulation (Barneche *et al.* 2014). A further, critical factor is the slow degradation rate of the transcriptional repressors, which mathematical analysis has long identified and experiments confirm as key to timing (Gerard *et al.* 2009, Ruoff *et al.* 2005). Regulated protein degradation is also crucial to plant circadian timing and provides one mechanism for environmental light signals to control the clock (Pudasaini *et al.* 2017).

### Functions of GI in the circadian clock

The *GIGANTEA* gene (*GI*) is central to protein regulation in the Arabidopsis clock. *gi* mutations delay flowering under long photoperiods that induce rapid flowering in wild-type plants Rédei et al. (1962), and also alter the pace of the circadian clock (Park et al, 1999; Fowler *et al*, 1999; Mizoguchi *et al*, 2005; Kevei *et al*, 2006; Martin-Tryon *et al*, 2007). GI affects the clock through interaction with the F-box proteins of the FKF1/LKP2/ZTL family, increasing the degradation of their target proteins, the evening-expressed circadian repressors PRR5 and TOC1 (Kiba *et al.* 2007, Mas *et al.* 2003). Degradation is mediated by the ARABIDOPSIS SKP1-LIKE (ASK) and CULLIN proteins, interacting with the F-box proteins to form a SKP-Cullin-F-box (SCF) ubiquitin ligase complex that targets proteins to the proteasome. GI RNA levels peak 8-10h after dawn (Dixon et al, 2011)(Fowler et al, 1999; David et al, 2006), before repression by the evening complex (composed of EARLY FLOWERING 3, ELF3, ELF4 and LUX (Nusinow et al, 2011). GI protein interacts with and is destabilized by ELF3 and COP1 (Yu et al, 2008), and is further partitioned between the nucleus and the cytoplasm. The constitutively-expressed, F-box protein ZEITLUPE (ZTL) and GI are mutually stabilizing. ZTL is therefore thought to enhance GI activity but also to sequester GI in the cytoplasm, whereas rhythmically-expressed GI confers rhythmicity upon ZTL protein levels (Kim *et al*, 2013d, 2013a, 2013c). Within the nucleus, the clock protein ELF4 interacts with and sequesters GI away from the promoter of the floral induction gene *CONSTANS (CO)*, contributing to rhythmic regulation of *CO* (Kim *et al*, 2013d).

### Role of GI in photoperiodic flowering induction

The rhythm of *CO* expression provides part of the timing function required to distinguish long, flower-inducing long photoperiods from less-inductive, short photoperiods. CONSTANS (CO) activates expression of *FLOWERING LOCUS T* (*FT*) (Samach *et al*, 2000). GI physically associates with the promoter regions of *CO* and *FT* (Sawa *et al.* 2007, Sawa & Kay 2011), and binds to transcriptional regulators (Kubota *et al.* 2017). Morning-expressed CYCLING DOF FACTORS (CDFs) *CDF1, CDF2, CDF3* and *CDF5* repress *CO* and *FT* transcription, preventing flowering in short days (Fornara *et al*, 2009; Song *et al*, 2012). FLAVIN-BINDING, KELCH REPEAT, F-BOX 1 (FKF1) is co-expressed with GI, binds both to GI and to CDF1-CDF5 under light conditions, and initiates their degradation by ubiquitylation (Imaizumi *et al*, 2005)(Fornara *et al.* 2009). Thus GI facilitates expression of *CO* and *FT* at the end of long photoperiods, by relieving CDF repression (Sawa *et al*, 2007; Fornara *et al*, 2009; Song *et al*, 2012).

### Broader roles of GI in metabolism and stress responses

*GI* has been linked to carbon metabolism, as it has a starch-excess phenotype (Eimert *et al.* 1995, Mugford *et al.* 2014) and is involved in a response of the circadian clock to sucrose (Dalchau *et al*, 2011). GI also plays a role in tolerance to a range of stress conditions, through sequestering the protein kinase SOS2. In high salt conditions, SOS2 is released to activate the Na^+^/H^+^ antiporter SOS1, leading to adaptation to higher salinity (Kim *et al*, 2013b). *GI* has been implicated in early flowering under drought conditions (Riboni *et al*, 2013). Finally, mutations in *GI* increase resistance to oxidative stress (Kurepa *et al*, 1998) and affect cold acclimation (Cao *et al*, 2005). GI’s biochemical mechanisms in most of these responses are unknown.

### Molecular mechanisms of GI function

GI’s role in the clock is mediated at the biochemical level by co-chaperone activity, which involves binding to HSP90 and appears to stabilize ZTL (Kim *et al*, 2011; Cha *et al*, 2017). This activity can affect other test substrates but its other native targets, if any, are unknown. As outlined above, GI’s known functions with the ZTL family and SOS2 are mediated by protein-protein interaction. Therefore, GI has been suggested to serve as a scaffold that orchestrates other protein interactions (Imaizumi & Kay, 2006), for example to provide chaperone activity (Cha *et al.* 2017).

Although such protein interactions are thought to mediate GI’s functions, these interactions have not been broadly tested. We therefore conducted interaction proteomics assays and obtained time-resolved data on potential direct and indirect partners of GI, over the daily timecourse. Here we discuss the abundance profiles and functions of candidate interactors and highlight a DOF protein, CDF6, validating its direct interactions and functional importance.

## Results

### Characterisation of a GI:3F6H line

We transformed Arabidopsis thaliana plants carrying the strong *gi-2* mutation (a deletion allele predicted to truncate ∼90% of GI protein, Fowler et al. 1999) with a construct to express 3xFlag-, 6xHis-tagged GI protein under the control of the CaMV 35S promoter (GI:3F6H). After isolating several positive transformants, we characterised one of the lines that expressed sufficient GI protein for tandem affinity purification (TAP) in a preliminary study (Figure S1). Analysis of specific gel bands by mass spectrometry (MS) identified GI and known interactors (Data S1). In addition, we tested expression of relevant genes in seedlings of this transgenic line using qPCR. The GI transcript was expressed constitutively, close to wild-type (WT) peak levels in the GI:3F6H line (Figure 1a). During the day, CO and FT mRNAs were higher in the GI:3F6H line than in the WT (Figures 1b, 1c), consistent with activation of these flowering-promoting genes by GI in the light (Sawa & Kay, 2011; Sawa et al, 2007). This was also reflected by slightly early flowering of the GI:3F6H plants relative to the WT, clearly rescuing the late flowering of the *gi-2* mutant parent (Figure 1d).

**Figure 1.**
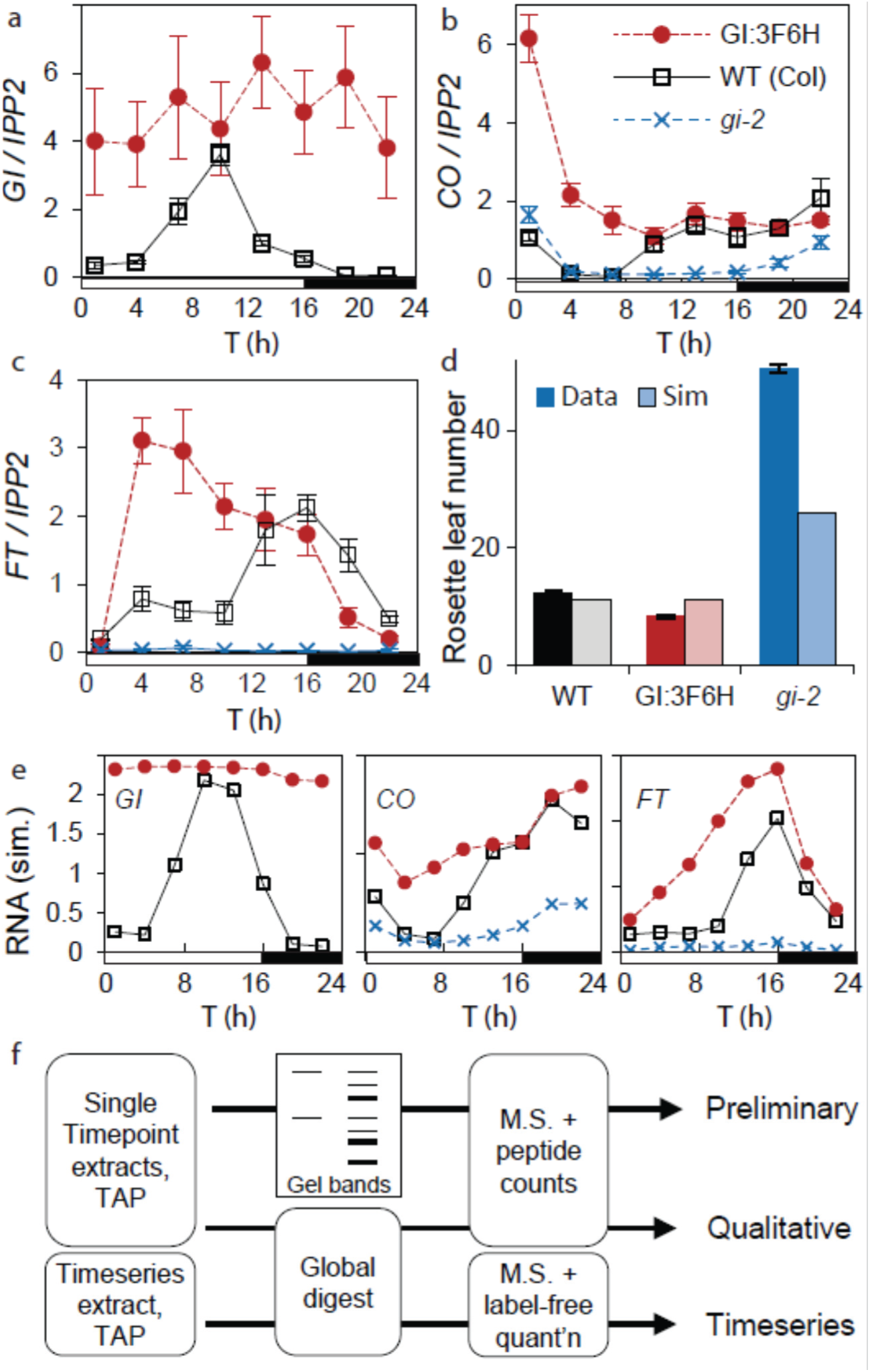
The GI:3F6H transgenic line restores flowering time. RNA expression was tested in samples of Arabidopsis plants of the Col WT, *gi-2* mutant and *gi-2* plants constitutively expressing the GI:3F6H fusion protein under long day conditions. qPCR assays detected GI (a), CO (b) or FT (c) RNA. Data are means of biological triplicates, normalized to an *IPP2* internal control; error bar, SEM. (d) Rosette leaf number was measured at flowering time in GI:3F6H, with WT and parental *gi-2* mutant controls, under long day conditions. Data (bold colours) are averages of 16 plants; error bar, SEM. The flowering experiment was simulated using the Framework Model v2 (FMv2; pale colours), as described in Supplementary Information. (e) *GI* transcription in the FMv2 was adjusted to match the *GI* RNA profile of GI:3F6H in (a); predicted expression profiles of *CO* and *FT* are shown (as in b, c). (f) Overview of proteomics studies using GI:3F6H.

Simulating the rescued mutant line in a mechanistic, mathematical model that includes photoperiodic flowering (Chew *et al.* 2017) predicted both molecular and flowering phenotypes (Figures 1d, 1e). The WT and *gi-2* simulations closely matched the RNA data, though this data set favoured morning (1-4h after dawn) expression of *CO* and *FT* compared to evening (13-16h) expression slightly more than the model, possibly reflecting a reduced *GI* RNA level at 13h. The model predicted elevated *CO* mRNA level at 1h in GI:3F6H plants but the observed level was even higher (Figure 1b). This *CO* peak induced more *FT* at 4h (Figure 1c) and earlier flowering in the data than the model.

The potential to identify further, GI-interacting proteins was tested in a qualitative proteomics study. Affinity purification of GI:3F6H (GI-TAP) was carried out with two different buffers (RIPA or SII) on GI:3F6H extract and a WT control for each buffer, followed by digestion and MS analysis (Figure 2b). 50 *Arabidopsis* proteins were identified by at least one peptide in each of the GI:3F6H samples and none in the controls (Table 1; full results in data S2). Detection of known direct interactors FKF1, ZTL and LKP2 validated the methodology, as well as indirect interactors ASK1A and ASK2, which are direct interactors of ZTL and FKF1 (Han et al, 2004; Kevei et al, 2006; Takahashi et al, 2004; Sawa et al, 2007). Among the remaining proteins on the list, half were abundant, ribosomal or cytoskeletal proteins or proteins presumably inaccessible to GI due to their annotated, chloroplast or mitochondrial location (but see Discussion). The remaining 17 proteins indicate the potential to identify previously undiscovered interactions, as discussed below.

**Figure 2.**
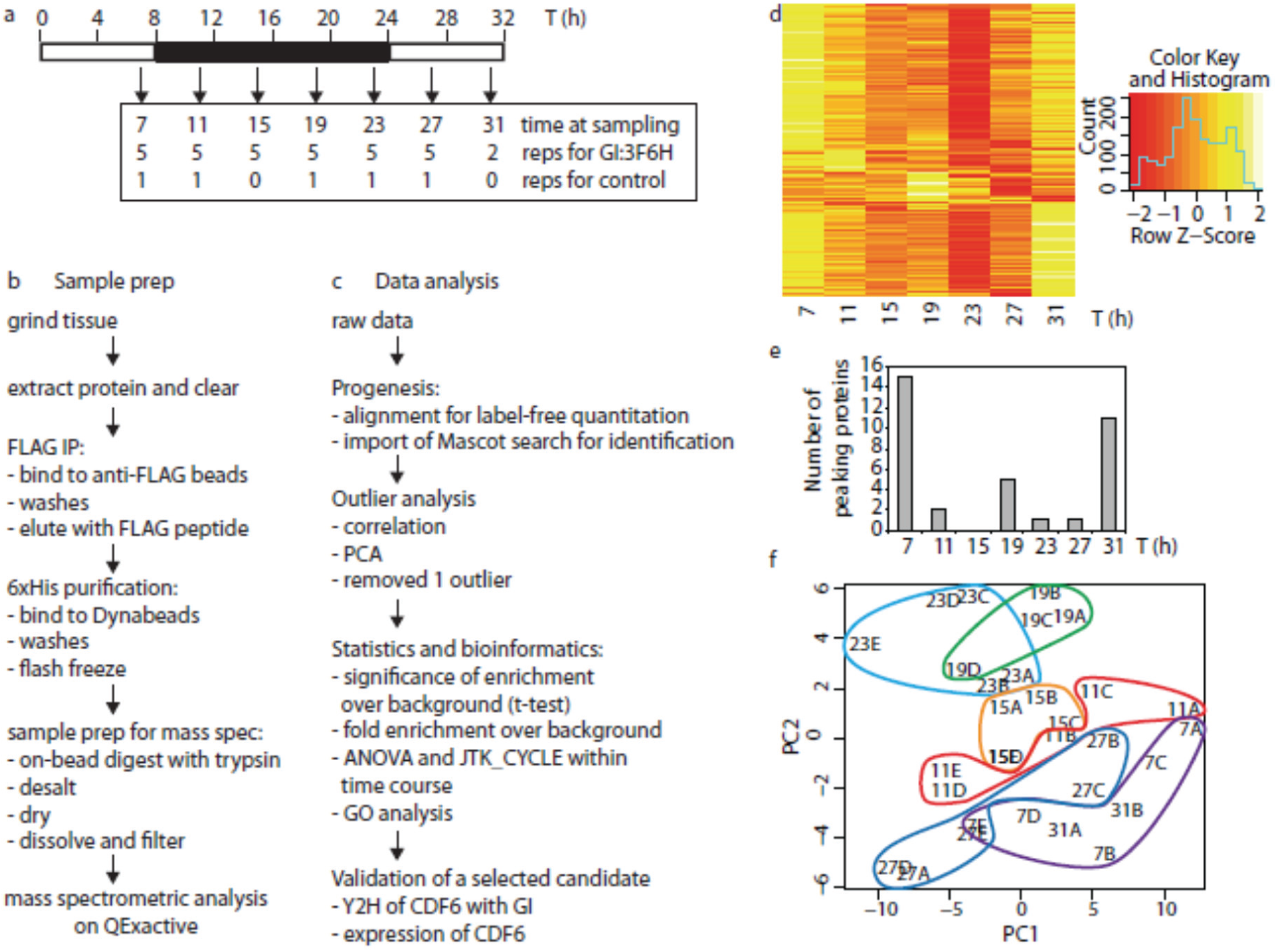
The GI-TAP timeseries study. (a) Samples from GI:3F6H and WT control plants grown in short day conditions were harvested at the indicated time points and replication. (b) Workflow for protein extraction, TAP using FLAG and His tags, and peptide preparation for mass spectrometry. (c) Workflow for label-free, quantitative data analysis, statistical and bioinformatics tests. (d) Heat map of protein abundance over time for 240 proteins with significant enrichment of at least two-fold. (e) Distribution of peak times, for 35 proteins shown in (d) with significant change over the GI-TAP time course (ANOVA p< 0.05 or JTK_CYCLE p< 0.05). (f) Principal component analysis separates GI-TAP timepoints (grouped by number and contour, with replicate letter) over components 1 and 2 (PC1, PC2). Note 31h timepoint replicates 7h.

**Table 1.**
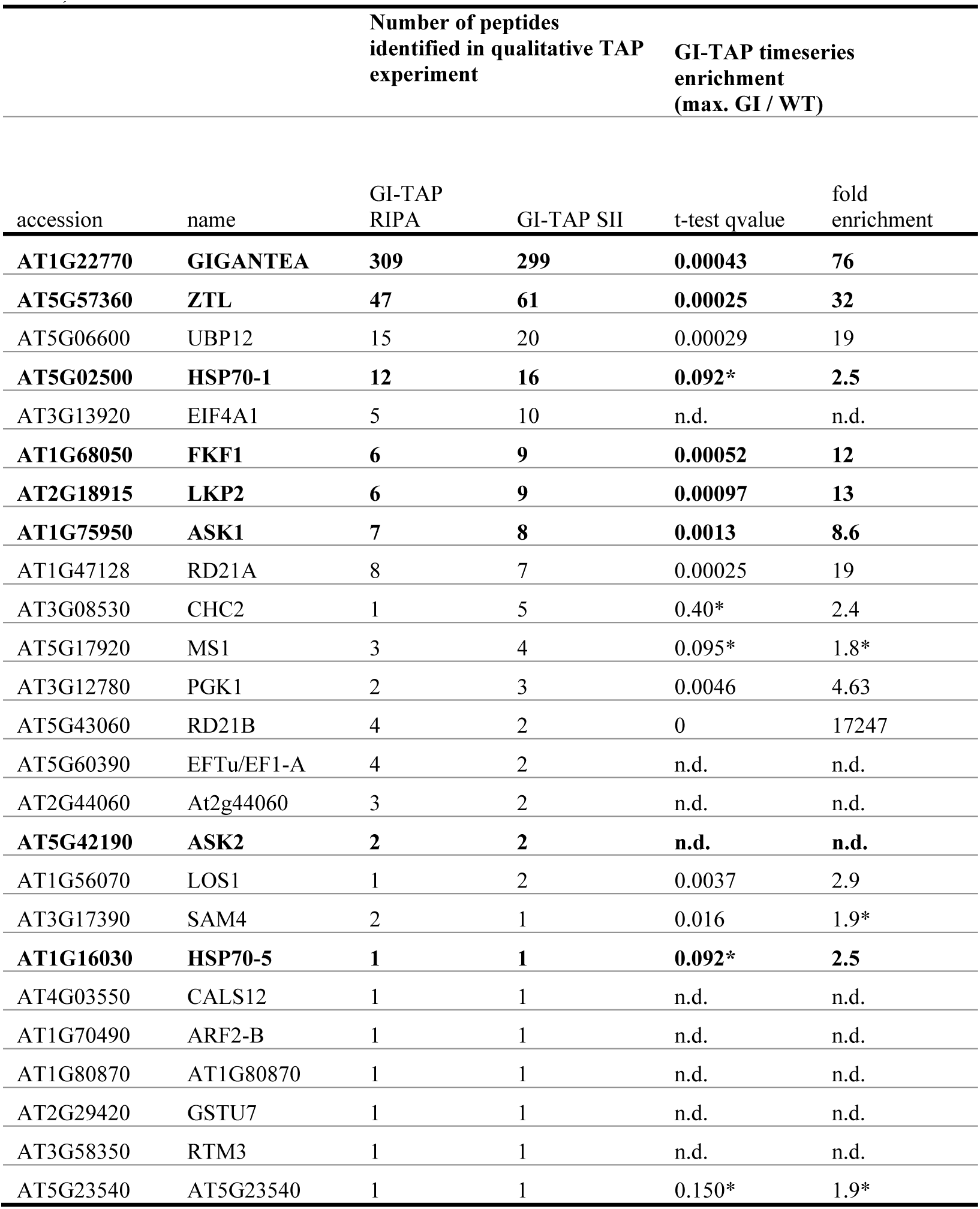
Candidate interacting proteins identified in the qualitative study. Control (WT tissue) and GI-TAP samples were extracted in RIPA or SII buffer. 25 Arabidopsis proteins were identified by at least one peptide in each GI-TAP sample and none in the controls, excluding likely contaminant proteins that are abundant (ribosome, cytoskeleton) or localized to other compartments than GI (chloroplast, mitochondria). The right–hand columns cross-reference the timeseries study (Table 3), with fold-enrichment and significance (q-value) of the maximum GI-TAP time point relative to the WT control. Bold: known direct or indirect interactors and homologues. *: below threshold in timeseries study. n.d., not detected.

### Candidate GI-interacting proteins from timeseries data

In order to obtain time-resolved interaction data, we applied the same GI-TAP method to plants grown in short-day conditions, at 6 timepoints in biological quintuplicate, with additional duplicate samples at time point 31h (replicating the 7h timepoint; Figure 2a). Extraction, TAP and sample preparation for mass spectrometry was carried out as for the qualitative analysis, using SII buffer (Figure 2b). In order to preserve low-abundance and indirect interactors, the experimental protocol did not remove all proteins that were bound non-specifically. Using the Mascot search engine to identify peptides, our choice of peptide score cutoff of 20 resulted in an FDR of 0.023. The mass spectrometry proteomics data have been deposited to the ProteomeXchange Consortium (<u>http://proteomecentral.proteomexchange.org</u>) via the PRIDE partner repository (Vizcaino *et al.* 2016) with the dataset identifier PXD006859 and DOI 10.6019/PXD006859. After identification and quantification of proteins (Figure 2c), one technical outlier was excluded from subsequent analysis (see Experimental Procedures; Figure S2). PCA of the remaining GI samples maximally separated the mid/late-night timepoints 19h and 23h from mid-day timepoints 7h and its replicate 31h (Figure 2e).

2336 peptides were detected in the time series study, from which 498 proteins were quantified (Table 2). The analytical methods also quantified the identified peptide peaks in WT samples, in contrast to the qualitative study. Therefore, the fold-enrichment of each protein in the GI-TAP was calculated relative to the average abundance in the control samples. Potential interacting proteins were identified as significantly enriched by t-test compared to the WT (excluding 8 proteins detected in only 1-2 WT samples, including RD21B, At5g43060), with a significance threshold adjusted for multiple testing (BH adjusted q-value < 0.05). Fold-enrichment threshold values were informed by the results for known interactors. The direct interactors FKF1, ZTL and LKP2 were more than ten-fold enriched over the control in the GI-TAP timeseries. Indirect interactors CUL1/CUL2 and GLN2 (Han et al, 2004; Song et al, 2014) were two- to three-fold more abundant at their peaks than the timeseries control and were not identified in the preliminary or qualitative studies. Hereafter we refer to significantly-enriched proteins with at least four-fold enrichment as highly enriched (136 proteins) and to proteins with two- to four-fold enrichment as weakly enriched (a further 104 proteins).

**Table 2.** Numbers of quantified and significant proteins in the GI-TAP timeseries. The raw data of the Progenesis output file (see Data S3) was used for statistics. * these numbers exclude proteins where t-test was impossible due to missing quantifications.

|  | 1 or more<br>unique<br>peptides | of these,<br>JTK_CYCLE<br>q value < 0.05 | 2 or more<br>unique<br>peptides | of these,<br>JTK_CYCLE<br>q value < 0.05 |
| --- | --- | --- | --- | --- |
| total identifications | 498 | 43 | 231 | 17 |
| significantly enriched (BH q-value < 0.05)* | 248 | 18 | 125 | 6 |
| and fold enrichment > 2 | 240 | 18 | 118 | 6 |
| and not abundant or in other location | 81 | 11 | 29 | 2 |
| and fold enrichment > 4 | 136 | 11 | 57 | 4 |
| and not abundant or in other location | 50 | 7 | 17 | 2 |

Among these 240 proteins, 25 changed significantly in abundance within the time series (assessed for any change by ANOVA) or 18 as assessed for rhythmic profiles (by JTK_CYCLE; Table 2 and data S3). Peak abundance for most of the 35 proteins with changing or rhythmic abundance occurred at 7h, or 24h later at 31h, with lowest abundance at 23h (Figure 2d). The abundance of the constitutively-expressed GI changed about two-fold over time (Figure 3a). The observed abundance changes for GI and ZTL closely matched the predicted protein profiles (Figure 3d) from simulation of GI:3F6H (as in Figure 1e). The change in GI abundance was not significant by ANOVA or JTK_cycle. Subsequent analysis used raw abundance rather than normalizing to remove the change in GI abundance; analysis of normalized data gave very similar results (data not shown and data S2). Our enriched protein sets should prioritise candidates for additional, direct or indirect interaction partners of GI, with greater scope than the smaller-scale studies.

**Figure 3.**
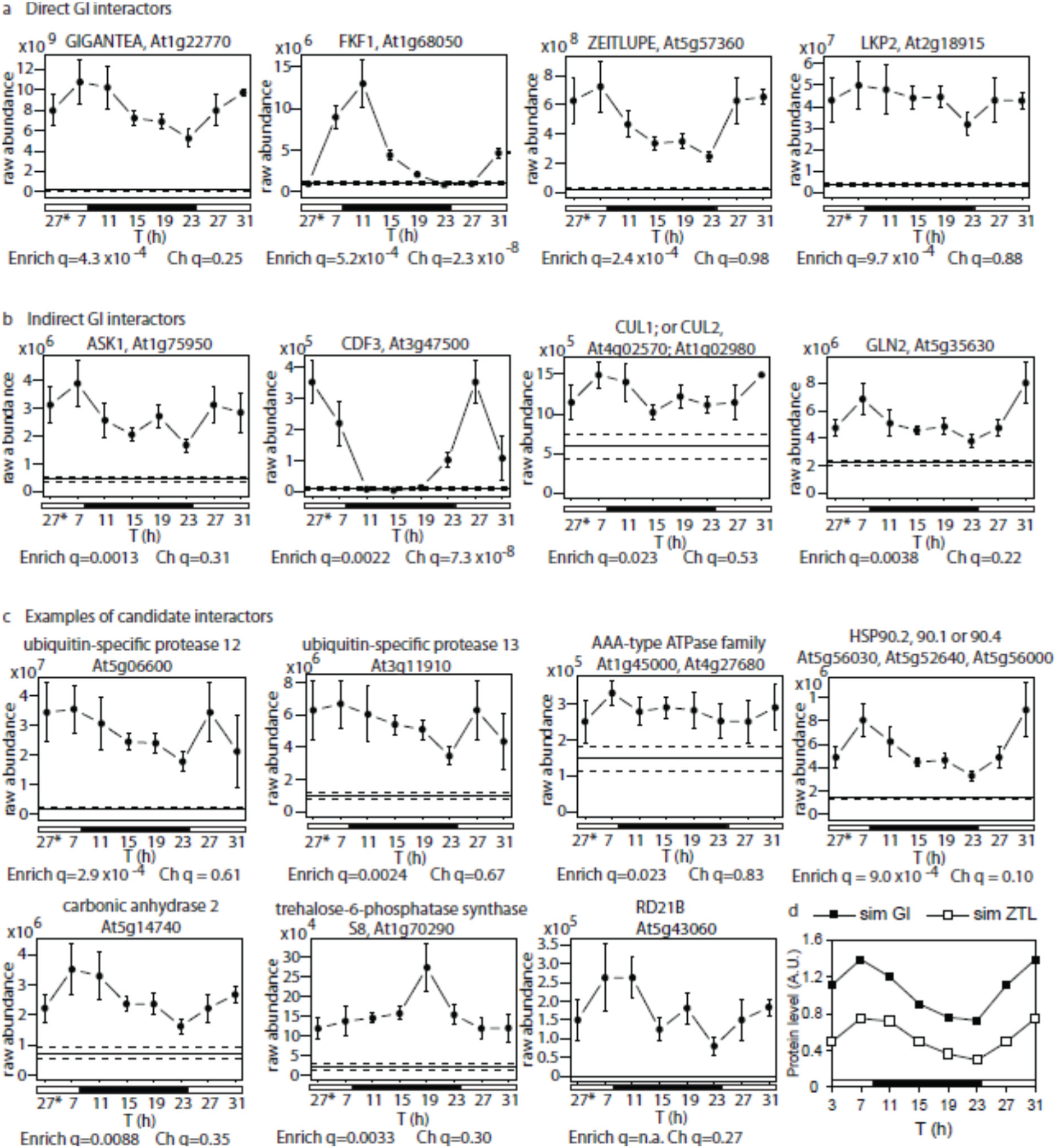
Diel profiles of GI and interacting proteins. Abundance of (a) GI protein and direct and (b) indirect interactors detected in the timeseries study showing that rhythmic profiles of FKF1 and CDF3 are atypical, along with (c) HSP90 and candidate (in)direct interactors. Multiple gene identifiers indicate related proteins were not distinguished by the three (CUL) or five (HSP90) peptides detected. GI-TAP samples, markers; error bar, SEM. Average of WT control, horizontal line, +/− SEM, dashed line. Time (T); * time point 27h is double-plotted to start the day in the morning. Significance of enrichment and temporal change are shown, as q-values of t-test comparing GI-TAP peak to WT (Enrich q) and of ANOVA within GI-TAP timeseries (Ch q). (d) Modelling the GI and ZTL protein timeseries, using the GI:3F6H model (Figure 1a) under short-day conditions, closely matches observations (panel a).

### Functional categorization of GI-TAP enriched proteins

According to a gene ontology (GO) search using the Panther tool (<u>http://pantherdb.org</u>, Thomas *et al*, 2003), the dataset contains 166 plastid proteins, of which 101 were weakly- or highly-enriched (data not shown). More detailed GO analysis in the topGO R package (Alexa *et al*, 2006), using all enriched proteins as foreground and all other identified proteins as background, revealed significant over-representation of plastid related terms (data S4, Figure S 3, appendix S1). GI protein is thought to be predominantly nuclear and cytosolic, so the analysis was repeated after excluding proteins annotated with predominantly chloroplast or mitochondrial localization. Abundant, cytoskeletal proteins were also excluded as likely non-specific interactors. The 35 remaining candidates are listed in Table 3, along with 3 related proteins. GO terms related to protein degradation were over-represented among the candidates (data S3), along with remaining, abundant protein classes such as glycolytic enzymes. The most significantly enriched term after ‘regulation of circadian rhythm’ was ‘ubiquitin-dependent protein catabolic process’ (GO:0006511; 7 enriched out of 14 detected proteins; p=0.0016).

**Table 3.**
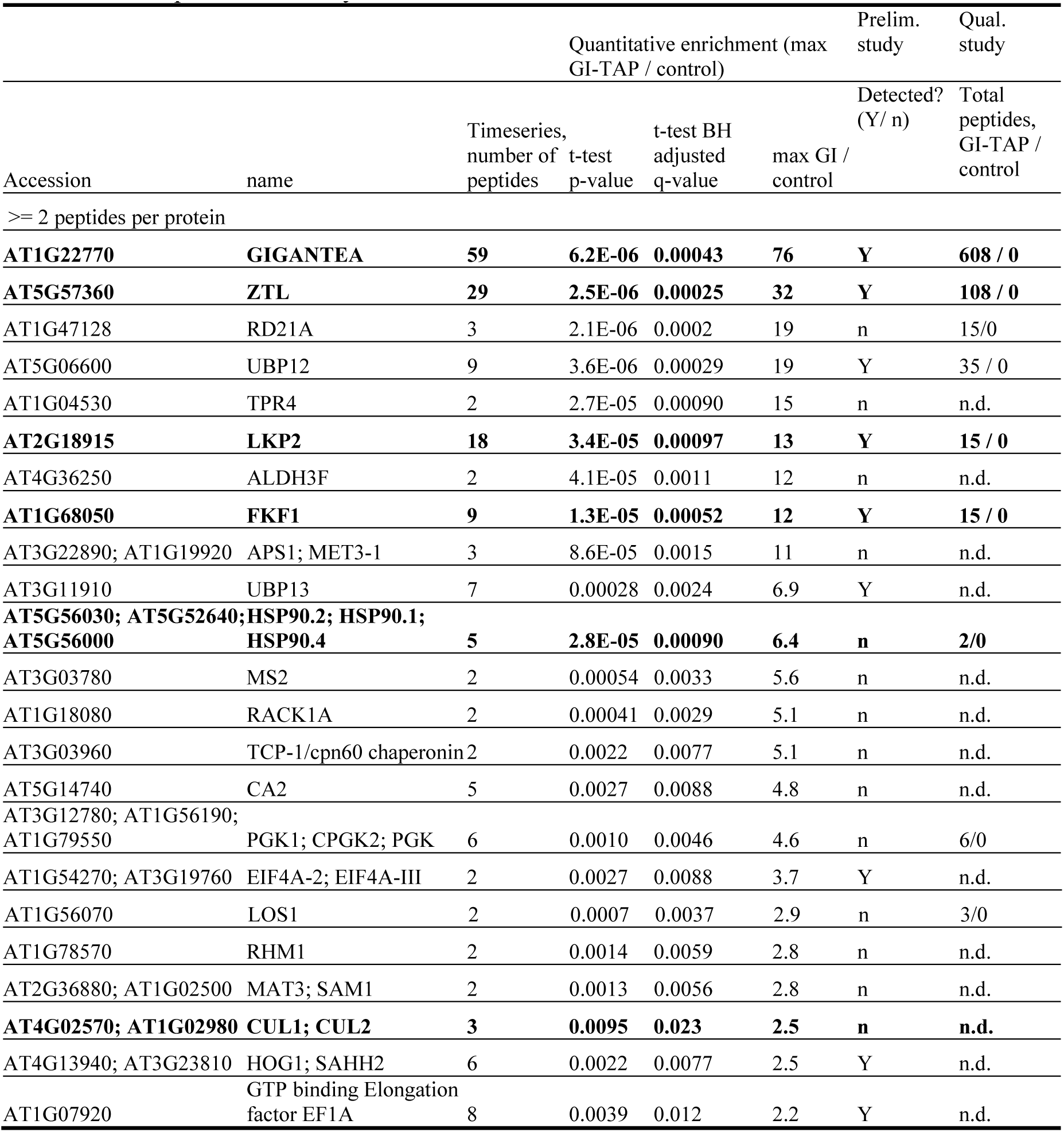
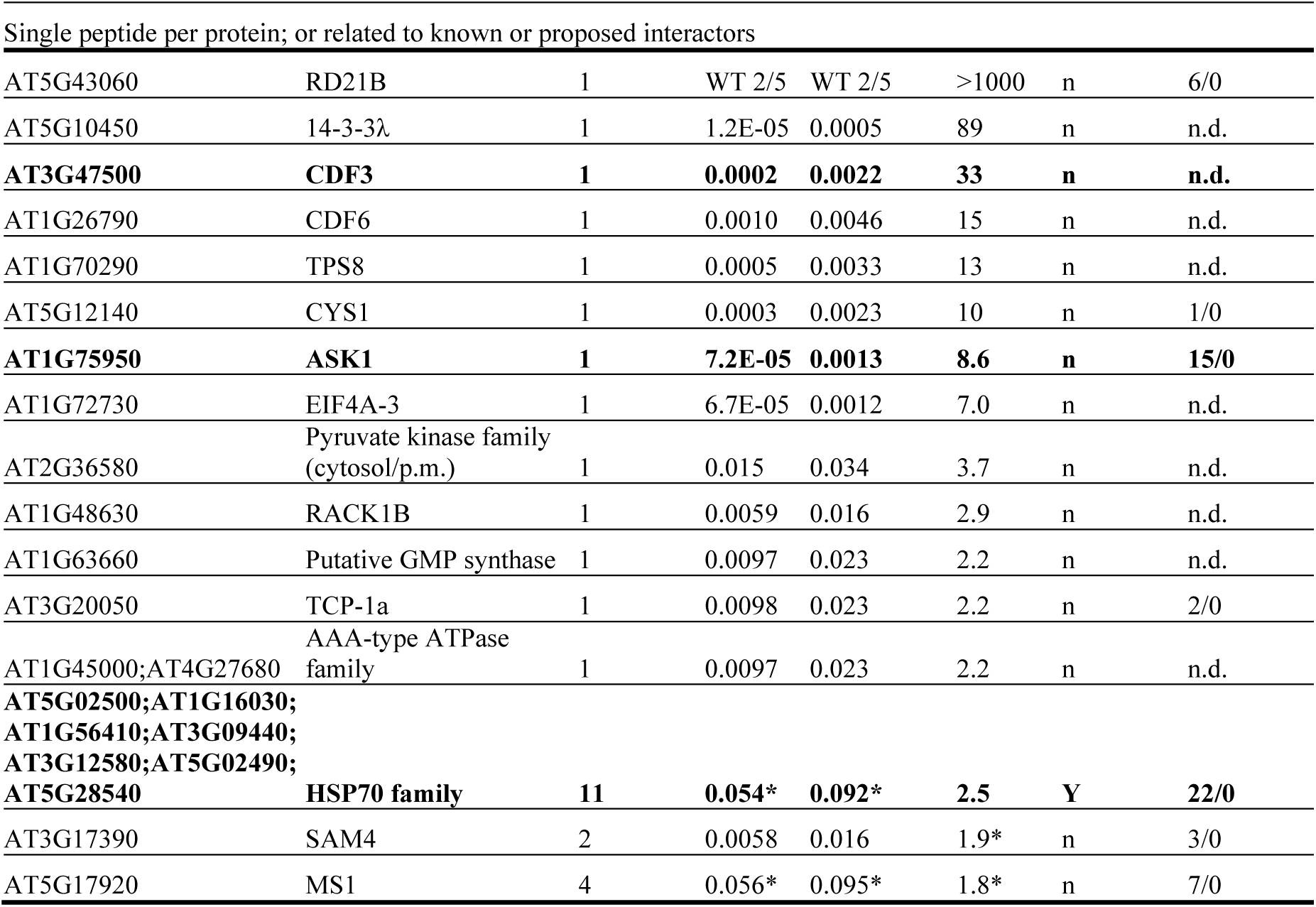
Known and candidate interactors of GI from GI-TAP time course experiment. Quantified proteins with two or more peptides that were significantly enriched (in t-test of maximum GI-TAP time point with WT control q-value < 0.05) by at least two-fold (max. GI-TAP / WT >2), ranked by fold enrichment. Only FKF1, CDF3 and CDF6 were rhythmic (JTK_cycle q-value < 0.05). Ribosomal or cytoskeletal proteins (abundant, likely unspecific interactors) and proteins in compartments putatively inaccessible to GI (chloroplasts, mitochondria) were omitted here (see Data S3). Where peptides matched very similar proteins, multiple accession numbers are shown. Bold type, known direct or indirect interactors. Detection in the preliminary study (Prelim.) is shown, and the sum of peptide numbers detected in the qualitative study (Qual.) in GI-TAP and WT (control). Selected proteins detected by single peptides are shown below, along with proteins suggested by other hypotheses (see Discussion) that were below thresholds in the timeseries (*) but were detected in the qualitative study.

### Rhythmic profiles of known (in)direct interactors

In contrast to the weakly rhythmic trend in abundance of the GI protein, known interactors showed contrasting profiles (Figure 3a). The direct interactors FKF1, ZTL and LKP2 showed temporal profiles consistent with their RNA expression patterns. Only FKF1 was strongly rhythmic, with a peak at 7-11h, resembling previous data (Fornara *et al.* 2009, Imaizumi *et al.* 2003). The ZTL profile paralleled GI, consistent with their mutual stabilization (Kim *et al*, 2013a) and closely matched by the prediction from the model simulation (Figure 3d). LKP2 abundance had a similar trend, consistent with arrhythmic RNA expression of *ZTL* and *LKP2*.

Four, established indirect interactors of GI were quantified (Figure 3b). CDF3 and GLUTAMINE SYNTHETASE 2 (GLN2) are client proteins of FKF1 (Song *et al*, 2014), whereas ZTL and LKP2 also interact with the core components of the SCF ubiquitin E3 ligase detected here, ASK1 and CUL1 and/or CUL2 (closely-related proteins that were not distinguished by the peptides detected). Of these, only CDF3 was scored rhythmic, with a peak at 27h and background levels between 11h and 19h. Therefore CDF3 level is in anti-phase with FKF1, in line with the degradation of CDFs by FKF1 (Fornara *et al.* 2009, Imaizumi *et al.* 2005). These results demonstrate the consistency of our data with published results. CDF3 was quantified using a single, individually-inspected peptide, indicating that such data should not be excluded from analysis.

The predicted functions of interactors include protein degradation and stabilization In addition to verifying the known indirect interactor CUL1, our analysis enriched other proteins involved in protein stability (Figure 3c). The stress-related proteases RD21A and RD21B were enriched in the timeseries and qualitative studies, as were ubiquitin-specific proteases (UBP) 12 and 13 (AT5G06600 and AT3G11910), AAA-type ATPase family proteins related to components of the 26S proteasome (At1g45000 and/or At4g27680) and a protease inhibitor, cystatin 1 (At5g12140). Several proteins annotated as being involved in protein stabilization were also significantly enriched by GI-TAP, such as Cpn60 chaperonin family proteins (At1g24510, At3g18190, At3g02530 and At3g03960) and at least one Hsp90 heat shock protein (At5g56030, At5g52640 and/or At5g56000, Figure 3c). Hsp70 family proteins were just below the enrichment cutoff (Table 3).

In contrast, neither additional F-box proteins nor other proteins involved in circadian timekeeping were identified as strong candidate interactors. The chloroplast RNA binding (CRB, At1g09340; Figure S3) is known to affect the amplitude of circadian clock and clock output genes (Hassidim *et al*, 2007) but the relevance of interaction with GI is unclear, given its localization (see Discussion). Multiple, metabolic enzymes and translation elongation or initiation factors were enriched in the GI-TAP timeseries. Among those that were not exclusively plastid localized (Table 3), were GTP-binding translation factors (At1g56070; At1g07920; At1g72730 and At1g54270 and/or At3g19760), two carbonic anhydrases (At1g70410 and At5g14740; Figure 3c and data S3), Phosphoglucomutase 2 and/or 3 (At1g23190, At1g70730; data S3), trehalose-6-phosphatase synthase 8 (TPS8; Figure 3c), a phosphofructokinase family protein (At1g20950), a GMP synthase homologue (At1g63660), a pyruvate kinase family protein (At2g36580) and 3 to 5 proteins of the methionine cycle, involved in regenerating S-adenosyl methionine (At3g03780; At2g36880/At1g02500; At4g13940/At3g23810) for methyltransferase reactions. At3g63410, At4g25080 and At3g63410 were the only enriched methyltransferases but all are plastid or mitochondrial-localised (Data S3). The candidate interactors suggest new clients and mechanisms of GI action related to those in other species (see Discussion), though their physiological significance awaits confirmation.

### CDF6 is a GI interactor that contributes to photoperiodic flowering

At1g26790 encodes a predicted DOF transcription factor that was up to 15-fold enriched around dawn (23h and 27h; Figure 4a). This protein was the most significantly rhythmic of the candidate interactors after FKF1 and CDF3 (BH-adjusted p value from JTK_cycle = 8 x 10^−6^). Its levels were the most anti-correlated with FKF1 levels among the highly-enriched proteins, followed by CDF3 (r = −0.65 and −0.40, respectively). We therefore validated the interaction and function of this protein, hereafter termed CYCLING DOF FACTOR 6 (CDF6). Yeast-2-hybrid (Y2H) assay experiments confirmed the interaction of full-length, N-terminal and C-terminal regions of GI with CDF6, as well as interaction of CDF6 with FKF1, ZTL and LKP2 (Figure 4b). *CDF6* transcript abundance was tested in plants transferred to constant light, revealing circadian regulation with a sharp peak around subjective dawn (Figure 4c), similar to *CDF1* transcript abundance (Seaton *et al*, 2015) and the profile of CDF6 in the GI-TAP timeseries (Figure 4a).

**Figure 4.**
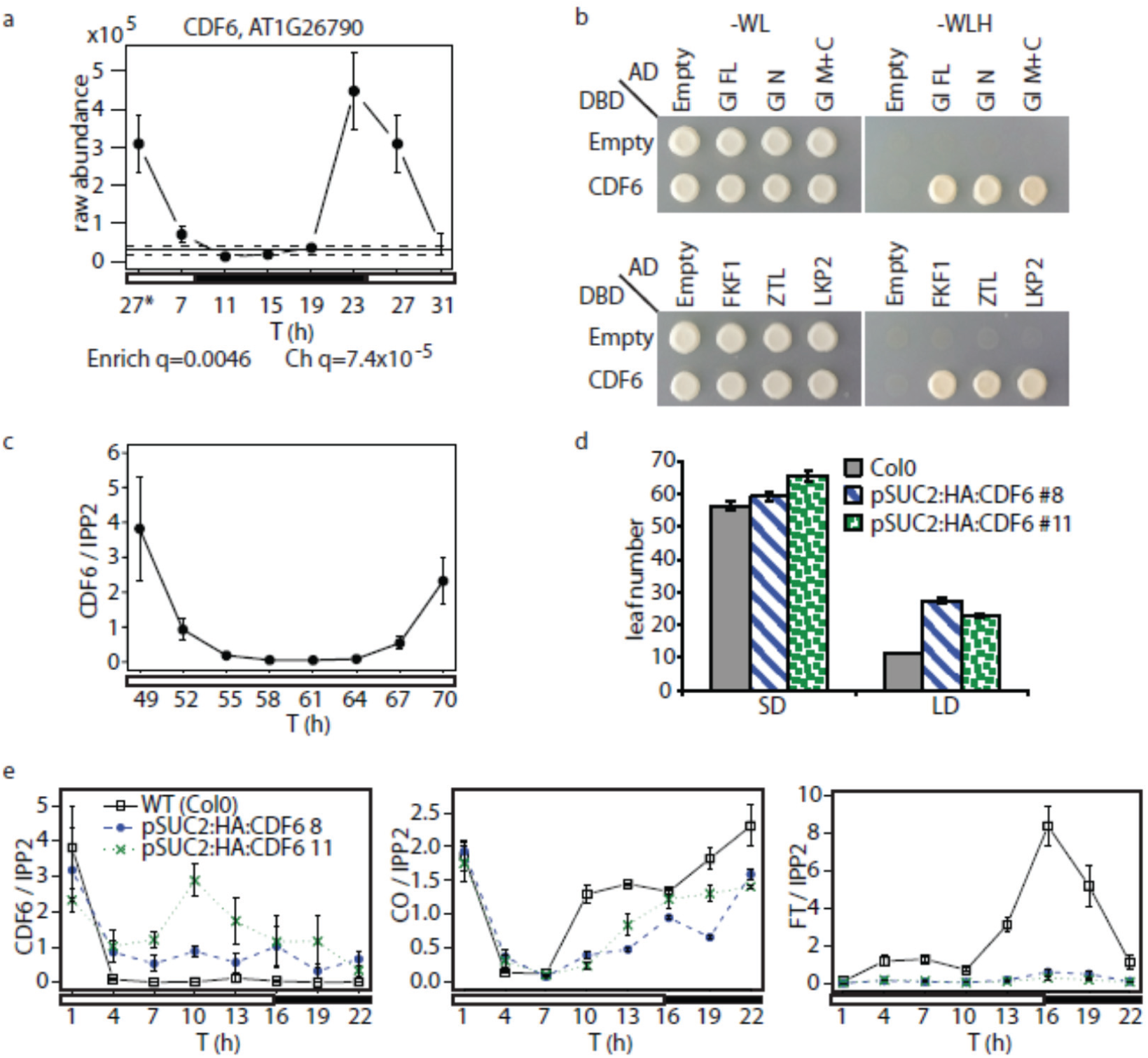
CDF6 interacts with GI and functions in photoperiodic flowering. (a) GI-interaction profile of CDF6 in the timeseries study is similar to CDF3 (Figure 3b). (b) Yeast two-hybrid assays validate interaction of CDF6 with full-length GI, N- and C-terminal domains of GI, as well as ZTL, FKF1 and LKP2. (c) Circadian expression profile of *CDF6* mRNA, in WT plants 3 days after transfer to constant light. (d) Constitutive *CDF6* expression delays flowering of transgenic SUC2-CDF6 lines more under long days, compared to WT control, than under short days. Each transgenic line differed significantly from WT, t-test p< 0.0001, except #8 in SD, not significant. (e) RNA expression profiles of *CDF6, CO* and *FT* were tested by qPCR in WT and the overexpressor lines, confirming that CDF6 suppresses evening *CO* and *FT* expression.

To test the potential function of CDF6 in the context of other members of the CDF gene family, transgenic plant lines were generated, in which CDF6 was expressed from the *SUC2* promoter that is active in phloem companion cells (Song *et al*, 2012). Two lines, pSUC2_HA:CDF6 #8 and #11, over-expressed the *CDF6* transcript at ZT4 and in later timepoints of a qPCR timeseries (Figure 4e), fluctuating around 20-60% of the WT peak level, whereas *CDF6* levels in WT were very low except at the ZT1 peak. Both transgenic lines flowered significantly later than WT under long photoperiods, with less effect under short photoperiods (Figure 4d). If CDF6 acts in a similar way to CDF1 (Song *et al*, 2012), we would expect it to inhibit transcription of both *CO* and *FT*. Indeed, *CO* RNA levels were reduced at 10h, 13h and at night in the transgenic plants compared to WT, and *FT* expression was reduced more than ten-fold at 4h and in later timepoints (Figure 4d). These results are consistent with CDF6 participating in the photoperiodic regulation of flowering, where CDF6 protein levels in WT are regulated through interaction with GI and its partners.

## Discussion

Proteostasis, the set of protein-metabolic processes, is expected to be critical for diel rhythms in general, because the removal of transcriptional repressor proteins controls the slow timing of circadian feedback circuits (Gerard *et al.* 2009, Ruoff *et al.* 2005). GIGANTEA not only indirectly mediates the degradation of transcriptional repressors (Kiba *et al.* 2007, Mas *et al.* 2003, Sawa *et al.* 2007a) but also has protein chaperone functions (Kim *et al*, 2011; Cha *et al*, 2017). Our studies identified further proteostatic proteins associated with GI, and suggested links to metabolic sensing, providing candidates for the unknown targets of GI’s proteostatic functions (Cha *et al*, 2017) and recalling previous data linking GI, metabolic inputs and biological timing.

### Technical requirements, known interactors and CDF6

Constitutive expression of tagged GI protein (GI:3FH6) in the *gi-2* mutant background rescued the mRNA expression and flowering time phenotypes of the mutant. GI protein tended to greater abundance during the day than during the night, in line with its light-dependent stabilization by ZTL (Kim *et al*, 2013a), with an evening peak time similar to native GI protein (David *et al*, 2006). The observed profile closely matched the prediction of a mechanistic clock model that was informed by diverse literature data (Pokhilko *et al.* 2012), indicating that the GI:3F6H protein conformed to the dynamic, light-responsive behaviour expected from previous results (Figure 3d).

The detection of known, indirect interactors of GI such as CUL1/2 among the weakly-enriched proteins but not in the qualitative studies validated the timeseries approach. Among the known, direct interactors of GI that were not detected in our studies, SVP, TEM1 and TEM2 (Sawa & Kay, 2011), COP1 and ELF3 (Yu *et al*, 2008), ELF4 (Kim *et al*, 2013d), CO (Song *et al*, 2014) and TCP4 (Kubota *et al*, 2017) are observed or expected to be largely or exclusively nuclear, while SPY (Tseng *et al*, 2004) and SOS2 (Kim *et al*, 2013b) are partly nuclear-localised. Analysis of nuclear preparations may be necessary to enrich for these and other, nuclear interactors. Rapid, whole-cell extraction was employed here to facilitate handling the larger sample numbers required to conduct the timeseries study in quintuplicate (Krahmer *et al.* 2015). Transcriptional regulators were nevertheless detected, including the known interactor CDF3 (At3g47500, Imaizumi *et al.* 2005) and its homologue CDF6 (At1g26790). Y2H assays validated the interaction of CDF6 with GI N- and C-terminal fragments, as well as with ZTL, FKF1 and LKP2 (Fornara *et al.* 2009). Functional overlap with other CDFs was confirmed, as constitutive *CDF6* expression inhibited *CO* and *FT* transcription and delayed flowering in a photoperiod-dependent manner (Figure 4d, e).

*CDF6* transcript expression in long days and constant light peaks around dawn, similar to *CDF1, CDF2, CDF3* and *CDF5* (Fornara *et al.* 2009, Imaizumi *et al.* 2005, Seaton *et al.* 2015). CDF6 interaction with GI was in antiphase to FKF1 interaction, consistent with CDF6 being largely or specifically degraded *via* this F-box protein. Our qualitative study and others conducted when GI normally accumulates (Huang *et al.* 2016a) coincide with peak FKF1 abundance, so would not have detected CDF6 or perhaps CDF3 (Figures 3b, 4a), confirming the utility of the timeseries approach. However only 35 proteins (15%) were enriched with a rhythmic profile, so the strong rhythms of FKF1 and the CDFs were uncommon. Rhythmic transcription of GI might confer rhythmicity on other partner proteins as it does for ZTL (Kim *et al.* 2013a), or the partners lack strong rhythmicity.

Conversely, the large size and proposed proteostasis functions of GI (discussed below) risk false-positive results. GI has not been found localized in or associated with the chloroplast but rather in the nucleus or cytoplasm (Huq *et al*, 2000; Kim *et al*, 2013a; Mizoguchi *et al*, 2005). The abundant, plastid-localised proteins enriched as interactors (Figure S3; Appendix S1) likely reflect unspecific binding, at least in the case of chloroplast-encoded proteins, which was an expected cost of detecting low-abundance and indirect interactors. Conservatively, we excluded cytoskeletal, ribosomal, mitochondrial and chloroplast proteins (see Methods; Data S3) from the candidate interactors (Tables 1 and 3). GI might in principle have a physiological role in the metabolism of proteins translated on cytosolic ribosomes, prior to compartmentalization, or of proteins translocated from other compartments to the cytosol for degradation (Svozil *et al.* 2014).

### Metabolic and nuclear functions of GI

The mechanisms that allow GI to link carbon metabolism and timing, via a long-term response of the circadian clock to sucrose (Dalchau *et al*, 2011) and the photoperiodic adjustment of the rate of starch biosynthesis (Mugford *et al*, 2014). The trehalose-6-phosphate (T6P) pathway mediates several such sugar responses (Ramon *et al*, 2009; Wahl *et al.* 2013). *TREHALOSE PHOSPHATE SYNTHASE 8* (*TPS8*) is a paralogue without known enzymatic activity but with diurnally-regulated expression, repressed by sucrose (Usadel *et al*, 2008). GI interaction with TPS8 was highly enriched and, unusually, peaked at ZT19 (Figure 3c), providing one of several possible mechanisms for GI to mediate between metabolism and biological timing. Strikingly, at least one of each enzyme in the cycle that regenerates the methyl donor S-adenosyl methionine (SAM; Figure S4) was enriched in the timeseries dataset, with 5-7 out of all 9 isoforms identified across our datasets (Table 3). The plastid-localised VTE3 was one of three enriched methyltransferases (Data S3). Interestingly in view of the Paraquat-sensitivity of *gi* mutants, VTE3 is expected to contribute *via* tocopherol synthesis to tolerance of reactive oxygen species, such as those generated by Paraquat treatment (Rastogi *et al.* 2014). The SESAME complex in yeast suggests an alternative hypothesis, as this chromatin modification complex links SAM synthetase with a histone methyltransferase and pyruvate kinase (Li *et al.* 2015). Indeed, a pyruvate kinase homologue (At2g36580) was also enriched in our data. The rhythmic abundance of GI might normally modulate methyltransferase activity, such as the rhythmic histone methylation at Arabidopsis clock genes (Barneche *et al.* 2014), or the broader chromatin methylation and flowering time affected by rice SAM synthetases (Li *et al.* 2011).

Several candidate interactors are shared with a previous study using ELF3 and ELF4 bait proteins (Huang *et al.* 2016a), which each interact with GI (Kim *et al.* 2013b, Yu *et al.* 2008): CRB has a known circadian role in plants (Hassidim *et al*, 2007); the 14-3-3λ protein (At5g10450) also interacts with known GI interactor SOS2 (Kim *et al*, 2013b)(Tan *et al.* 2016) and related 14-3-3 proteins interact with GI (Chang *et al.* 2009) and the Arabidopsis S-adenosylmethionine synthetases (Shin *et al.* 2011); RACK1A (At1g18080) is a promiscuously-interacting protein with several reported physiological roles in plants (Islas-Flores *et al.* 2015). Its homologue RACK1B (At1g48630) was also weakly enriched (Table 3). Mammalian RACK1 affects the circadian clock through the interacting transcription factor BMAL1 (Robles *et al.* 2010), and contributes to degradation of its paralogue hypoxia-induced factor HIF1a. HIF1a protein regulation is mediated *via* Hsp90 and ubiquitin-specific proteases (reviewed in Wilkins *et al.* 2016): their Arabidopsis homologues were highly enriched in our GI-TAP datasets.

### GI interactors involved in protein metabolism

Protein degradation of clock-relevant, transcriptional repressors was the first biochemical function supported for GI, acting as a scaffold for F-box proteins, though GI’s co-chaperone function is now also implicated (Cha *et al*, 2017; Kim *et al*, 2007). No further F-box proteins or other ubiquitin E3 ligases were identified here, suggesting that GI mediates further physiological roles through different biochemical mechanisms. Involvement in translational regulation was suggested by the enrichment of translation-associated proteins, including the LOS1 protein (Tables 1, 3) implicated in cold stress (Guo *et al.* 2002). Chaperone proteins are typical, non-specific contaminants of affinity purification studies but direct, physiologically-relevant binding of HSP90 with GI has been demonstrated (Cha *et al.* 2017). HSP90 isoform(s) were highly enriched and weakly rhythmic in our timeseries (Figure 3c). TPR4, which encodes a tetratricopeptide repeat (TPR) protein with potential to interact with Hsp90/Hsp70 as a co-chaperone (Prasad *et al.* 2010) was also strongly enriched (Table 3). The Hsp70 family proteins that might function with GI and HSP90 (Cha *et al.* 2017) were below the significance threshold in the timeseries study (Table 3) but one (At5g02500) was the fourth most-enriched protein in the qualitative study (Table 1).

In contrast, several other proteins involved in proteostasis were highly and reproducibly enriched. TCP-1/cpn60 chaperonin family proteins (At3g03960; At3g20050) that can facilitate intercellular trafficking of transcription factors (Xu *et al.* 2011) were detected in the timeseries and qualitative studies (Tables 1, 3). Granulin-repeat cysteine proteases RD21A and RD21B (Figure 3c) were also repeatedly detected. These are among the major protease activities of the leaf and are inducible by drought stress and senescence (Koizumi et al, 1993; Gu *et al.* 2012). Their functional relationship to the other stress-related GI interactors SOS2 and 14-3-3λ is unclear. RD21A was previously identified as an interactor of three transcription factors (Huang *et al.* 2016a, Huang *et al.* 2016b, Wang *et al.* 2013), two of which interact with 14-3-3λ, and also with the final, proteostatic interactors of GI, the ubiquitin-specific proteases UBP12 and UBP13.

UBP12 and UBP13 were highly enriched in the timeseries and were also detected in the preliminary and/or qualitative studies. Their de-ubiquitination activity potentially counteracts protein degradation, for example of Arabidopsis MYC2 (Jeong *et al.* 2017), or monoubiquitination, for example of histone H2A (Derkacheva *et al.* 2016). UBP12 and UBP13 are already known to affect the Arabidopsis circadian clock, act upstream of *GI* and *CO* in the same photoperiodic flowering time pathway (Cui *et al*, 2013), and are recruited to chromatin in association with the histone methylation complex PRC2 (Derkacheva *et al.* 2016). In a final connection, histone de-ubiquitylation by USP7, the Drosophila homologue of UBP12/UBP13, is allosterically controlled by its interaction with a GMP synthetase (van der Knaap *et al.* 2005). An Arabidopsis homologue (At1g63660) of this enzyme was also enriched in the timeseries data (Table 3). Thus the GI-TAP results not only identified a rhythmic CDF protein implicated in the canonical, flowering pathway but also suggest that plants share the intriguing links among proteostatic functions, metabolic enzymes and chromatin modification that are now being identified in other organisms (van der Knaap and Verrijzer 2016). These provide a novel set of hypotheses on the biochemical mechanisms of flowering regulation and of further physiological effects of GI.

## Experimental Procedures

### Generation of plant materials

To generate plants with epitope-tagged GI protein, *gi-2* mutant *Arabidopsis thaliana* plants were transformed with a construct expressing C-terminally 3xFLAG-6His tagged GIGANTEA protein (GI:3F6H). The full-length GI cDNA without the stop codon was amplified and inserted into pENTR/D-TOPO vector (Invitrogen). After sequence verification, the *GI* cDNA was transferred into the pB7HFC vector, designed for in-frame epitope fusion (Huang et al., 2016) by a Gateway cloning reaction (Invitrogen). pB7HFC-GI-3F6H was introduced to the *gi-2* mutant by Agrobacterium-mediated transformation. Transgenic plants that rescued the *gi-2* phenotype were selected, and the expression of the GI:3F6H transgene was verified by Western blotting (as in Figure S1; Methods S1). Samples for the preliminary and qualitative GI-TAP studies were prepared as described in (Song et al., 2014).

To generate *SUC2:HA-CDF6* plants, the *CDF6* CDS was PCR-amplified using cDNA derived from long-day grown plants as a template, and cloned into pENTR D-TOPO (Invitrogen), to form pENTR HA-CDF6. 2.3 kbp of the *SUC2* 5' upstream promoter region was amplified and cloned into the pENTR 5'-TOPO vector (Invitrogen), to form pENTR 5' SUC2. Using a sequential LR clonase II reaction (Invitrogen), we integrated the pENTR 5' SUC2, pENTR HA-CDF6 into the R4pGWB501 vector (Nakagawa *et al.* 2008), to form *SUC2:HA-CDF6*. After confirming the sequence, this vector was transformed into Col-0 wild-type plants using by *Agrobacterium*-mediated transformation. Transgenic plants were selected based on the expression level of *CDF6* RNA.

### Plant growth conditions

For flowering time experiments, seeds were sown and stratified at 4 °C for 3 days on soil (Sunshine Mix #4; Sun Gro Horticulture), containing Osmocote Classic time-release fertilizer (Scotts, Marysville) and Systemic Granules: Insect Control (Bionide, Oriskany). Growth was at 22 °C under long-day conditions (16h light, full-spectrum white fluorescent light bulbs (F017/950/24′′ Octron; Osram Sylvania, 70-80 μmol·m^−2^·s^−1^). Flowering time was measured as the mean number of rosette leaves, for at least 16 plants per genotype, ± the standard error of the mean (SEM).

For qPCR analysis, 10-day old seedlings were grown on 1× Linsmaier and Skoog (LS) media (Caisson), supplemented with 3% (w/v) sucrose and 0.8% (w/v) agar, under long-day conditions at 22°C in growth chambers (CU-36L5; Percival Scientific; lighting conditions as for flowering time) and harvested at 3-h intervals from 1 h after dawn.

For the proteomics timeseries, GI:3F6H and Col-0 WT seeds were surface-sterilized for 10 min with 30% thin bleach, 0.01% TritonX-100, followed by 4 washes with sterile water. After cold-treatment at 4°C for 5 days, seeds were grown on plates (2.15g/l Murashige & Skoog medium Basal Salt Mixture (Duchefa Biochemie), pH 5.8 (adjusted with KOH) and 12g/l agar (Sigma)) in Percival incubators (CLF Climatics) for 17 days at 85µmol m^−2^s^−1^ (full-spectrum white fluorescent bulbs) and 21°C in short day conditions (8h light). Seedlings were transferred to soil, for 20 days in the same conditions with a light intensity of 110 µmol m^−2^s^−1^. Starting at 7h after dawn, 80 rosettes without roots were harvested for each sample, at timepoints and replicates shown in Figure 2a and flash-frozen in liquid nitrogen. Dim green safelight was used during darkness.

### Quantitative PCR (qPCR) analysis

Fresh seedlings were ground into powder with a mortar and pestle with liquid nitrogen, and total RNA was isolated by using an illustra RNAspin Mini kit (GE Healthcare) according to the manufacturer’s instructions. Two µg of total RNA was reverse-transcribed using the iScript cDNA synthesis kit (Bio-Rad) according to manufacturer’s instructions. cDNA was diluted to 5 times with water, and 2 μl was used as a template for quantitative PCR (qPCR) analysis using primers shown in Table S1. Isopentenyl pyrophosphate/dimethylallyl pyrophosphate isomerase (IPP2) was used as an internal control for normalization. The average value from WT was set to 1.0 to calculate the relative expression of other lines. To amplify *CO* and *CDF6*, three-step PCR cycling program was used: 1 min at 95 °C, followed by 40–50 cycles of 10 s at 95 °C, melting temperatures for 15 or 20 s, and 72 °C extension for 15 s. To amplify *GI*, *FT*, and *IPP2*, a two-step PCR cycling program was used: 1 min at 95 °C, followed by 40–50 cycles of 10 s at 95 °C and 20 s at 60 °C. Data show the average of three biological replicates with SEM; each measurement had two technical replicates.

### Protein Extraction and tandem affinity purification (TAP) procedure

All steps in the protein extraction, TAP and preparation for mass spectrometry were carried out in random sample order in order to avoid bias due to order of processing. Frozen plant tissue was ground to a fine powder in a liquid nitrogen and dry ice-cooled mortar and processed essentially as described (Song et al., 2014). Detailed procedures are described in Supporting Experimental Procedures (Methods S2).

### Protein digestion and mass spectrometric analysis

Preparation of samples for mass spectrometry for the qualitative and timeseries studies analysis used an on-bead digest, prior to mass spectrometric analysis. Detailed procedures are described in Supporting Experimental Procedures (Methods S3).

### Proteomics data analysis and bioinformatics

For the qualitative study, database searches were performed using Comet (Eng *et al.* 2013), searching against the Uniprot Arabidopsis protein sequence database, and using Percolator (Matrix Science, Boston, USA) with a q-value cutoff of 0.01. Cysteine residue masses were considered statically modified by iodoacetamide, and methionine dynamically modified by a single oxidation. Precursor mass tolerance was 10 ppm, and product ion tolerance 0.5 Da. The principle of parsimony was used for protein inference, and at least two unique peptides were required for each identified protein.

The timeseries data were analysed using the commercial Progenesis LC-MS software (version 4.1.4924.40586; Nonlinear Dynamics, Newcastle) for label-free quantitation. Raw files were imported into a label-free analysis experiment, chromatograms were subjected to automatic alignment and peak picking. Only charges 2+, 3+ and 4+ and data from 25 to 75 minutes of the runs were chosen for analysis. The exported file of MS/MS spectra was uploaded on the Mascot website (version 2.4) and a search was carried out with the following parameters: database Arabidopsis_1rep (version 20110103), trypsin as enzyme, allowing up to two missed cleavages, carbamidomethyl (C) as a fixed modification, Oxidation (M), Phospho (ST) and Phospho (Y), as variable modifications, a peptide tolerance of 10ppm, and MS/MS tolerance of 0.05Da, peptide charges 2+, 3+ and 4+, on a QExactive instrument (Thermo), and with decoy search to determine false discovery rate (FDR). For export, an ion-cutoff of 20 was chosen. Protein and peptide measurements were exported to identify technical outliers, using correlation analysis and principal component analysis (PCA) implemented by an R script (data S5). The average Pearson correlation coefficient of each GI-TAP replicate with the other replicates of the same time point was above 95% for all GI:3F6H samples apart from sample 19E (Figure S2), which was also clearly separated from all other samples by PCA and was therefore discarded.

A custom R script performed further statistical analysis and plotting (data S7). We used a t-test to determine for each protein, whether the maximum GI-TAP time point (omitting the 31h time point) is significantly different from the WT control average using q-values (Benjamini-Hochberg (BH) corrected p-values). ‘Fold enrichment’ is the ratio of the highest GI-TAP timepoint to the WT control average gives. To assess temporal changes, ANOVA was performed on arcsinh-transformed GI-TAP data, including the ZT31 timepoint. To assess rhythmicity, we used the JTK_CYCLE tool (Hughes *et al*, 2010) to analyse periods of 22-26h. The summary heatmap (Figure 2d) used the heatmap.2 function of the pvclust v2.0 R package (Suzuki & Shimodaira, 2015). Gene ontology (GO) analysis was performed using topGO (http://bioconductor.org/biocLite.R, version: 2.16.0, (Alexa & Rahnenführer, 2010), using a node size of 3, as described by (Krahmer et al, 2015; Data S6). Biological context was provided by subcellular locations annotated in the SUBA resource (Hooper *et al.* 2017) and interaction data in the BioGrid (Chatr-aryamontri *et al.* 2017).

### Yeast-2-Hybrid assay

Full-length *CDF6* coding sequence was PCR-amplified using cDNA as template with primers shown in Table S1, cloned into pENTR D-TOPO (Invitrogen) and sequence-verified. The plasmid cassette was transferred to pAS-GW, a gateway compatible bait vector (Nakayama *et al.* 2002) using LR clonase II (Invitrogen). The GI-N, GI-C, FKF1, LKP2, and ZTL clones used in this analysis were described previously; GI-N and GI-C (Sawa *et al.* 2007b), and FKF1, LKP2, and ZTL (Imaizumi *et al.* 2005). Yeast strains Y187 and AH109 were transformed with prey and bait vectors, respectively using the standard yeast transformation protocol (Clontech). After colonies formed on –W or –L containing media, three independent colonies were grown together, and then mated against their corresponding pairs for 3 days on YPDA media. After mating, yeast colonies were transferred onto –WL media. After checking for mating confirmation, yeast sectors were retransferred at the same time onto –WL and – WLH media. The experiments were repeated several times with the same results.

### Accession numbers

AT5G43060, AT5G57360, AT1G22770, AT1G68050, AT1G56500, AT1G04530,

AT5G02940, AT5G06600, AT1G23410, AT3G09790, AT1G47128, AT4G01050,

AT5G48300, AT2G18915, AT3G56940, AT1G60780, AT5G09810, AT2G42170,

AT3G12110, AT1G06950, AT1G15930, AT4G35250, AT5G01220, AT3G11910,

AT5G01920, AT1G56070, AT1G31330, AT1G05190, ATCG00270, AT5G56030,

AT5G52640, AT5G56000, AT4G04640, AT1G55670, AT5G47210, AT3G62910,

AT1G75950, AT2G39800, AT3G63410, ATCG00020, ATCG00120, AT1G04170,

AT5G03880, AT1G48120, AT1G67700, AT5G43970, AT5G54770, AT1G08520,

AT5G26710, AT3G48140, AT1G22530, AT2G17630, AT1G14810, AT3G22890,

AT1G19920, AT4G20360, ATCG00670, AT3G47500, AT2G40660, AT5G51110,

AT3G03780, AT1G61520, AT5G35630, AT2G29550, AT1G07410, AT1G01200,

AT3G44310, AT3G44320, AT5G22300, AT4G25630, AT1G29880, AT4G27440,

AT5G54190, AT4G10340, AT5G50920, AT2G25140, AT3G48870, AT2G39770,

AT1G22410, AT4G39980, AT4G11150, ATCG00780, AT2G38750, AT4G25080,

AT1G49670, AT5G12250, ATCG00130, AT1G09180, AT4G37800, ATCG00480,

AT5G08670, AT4G22890, AT3G03960, AT2G18020, AT3G51190, AT4G36130,

AT5G46290, AT5G44340, AT3G25920, AT3G60300, AT1G20010, AT1G75780,

AT1G16410, AT1G20620, ATCG00680, AT1G10630, AT1G04480, AT4G01690,

AT4G39520, AT1G26790, AT3G02530, AT1G18080, AT3G11710, AT3G12780,

AT1G56190, AT1G79550, AT2G39730, AT4G31580, AT5G52840, AT1G78580,

AT5G28060, AT3G04920, AT1G64880, AT3G10260, AT1G26630, AT3G44110,

AT5G22060, AT1G50370, AT3G02090, AT3G08940, AT4G10320, AT3G23145,

AT3G22960, AT1G79040, AT1G19880, AT1G48630, AT2G45710, AT2G07698,

ATMG01190, AT1G49760, AT1G74470, AT1G09130, AT5G01530, AT4G18240,

AT3G02320, AT5G23060, AT5G19770, AT1G64740, AT3G02780, AT2G37990,

AT2G17130, AT1G78900, AT5G38660, AT3G20050, AT5G14740, AT2G19730,

AT5G52040, AT4G25500, AT1G72370, AT3G04770, AT5G10450, AT2G21580,

AT5G35980, AT3G06650, AT4G02570, AT1G02980, AT5G20010, AT1G69830,

AT5G60100, AT5G65730, AT5G63150, AT1G07770, AT2G39590, AT3G46040,

AT5G06850, AT3G02080, AT4G05180, AT3G05060, AT3G46780, AT1G04820,

AT4G23920, AT4G10960, AT5G02500, AT1G16030, AT1G56410, AT3G09440,

AT3G12580, AT5G02490, AT5G28540, AT3G29360, AT2G47650, AT3G18190,

AT5G54600, AT5G40270, AT5G40290, AT4G20890, AT1G07660, AT5G67500,

AT5G03290, AT3G09810, AT3G09880, AT1G13460, AT3G21650, AT4G13930,

AT2G21160, AT2G34430, ATCG00280, AT2G36160, AT3G11510, AT2G05070,

AT2G36880, AT1G02500, AT5G47010, AT1G62750, AT3G60750, AT2G45290,

AT1G44446, AT1G59870, AT1G54270, AT3G19760, AT5G42270, AT1G50250,

AT3G53420, AT2G37170, AT5G60660, AT5G11710, AT1G22780, AT3G04120,

AT1G13440, ATCG00830, AT1G30380, AT1G72730, AT1G29930, AT1G29910,

AT2G47710, AT3G10670, AT1G11220, AT2G16360, AT4G34670, AT3G06400,

AT3G44300, AT3G01500, AT1G08730, AT5G45530, AT4G27700, AT5G02870,

AT3G03950, AT2G28000, AT3G14420, AT4G18360, AT5G59950, AT1G02150,

AT3G32930, AT2G30860, AT2G30870, AT1G75750, AT5G23860, AT1G02560,

AT3G51600, AT3G44890, AT3G08590, AT1G09780, AT4G31420, AT5G62690,

AT3G45140, AT1G45000, AT4G27680, AT5G64040, AT1G80410, AT5G61670,

AT1G71500, AT4G21960, AT5G63620, AT4G35630, AT1G07790, AT2G37470,

AT5G35200, AT1G68830, AT1G42970, AT3G42050, AT4G11010, ATCG00330,

AT1G54410, AT2G26250, AT1G15690, ATCG00490, AT2G07732, AT1G48610,

AT1G80480, AT5G61220, AT1G49970, AT1G58380, ATCG00470, AT5G13650,

AT5G19430, AT4G15560, AT3G21500, ATCG00800, AT3G24830, AT1G48350,

AT1G12900, AT1G55490, AT5G56500, AT4G15000, AT2G32220, AT3G22230,

AT1G37130, AT5G30510, AT1G15820, AT1G24510, AT5G40950, AT3G58140,

AT1G14510, AT4G02520, AT4G03020, AT1G49780, AT1G18170, AT1G74050,

AT1G18540, AT3G01310, AT3G48730, AT1G08880, AT1G16300, AT4G16340,

AT5G12470, AT2G36460, AT3G52930, AT3G62120, AT1G54520, AT2G37190,

AT5G26000, AT5G20290, AT1G20330, AT1G32220, AT2G33040, AT4G38970,

AT4G37930, AT5G26780, AT2G43030, AT2G36620, AT1G74730, AT1G56200,

AT1G16720, AT1G23310, AT4G36250, AT3G60770, AT5G12140, AT2G04030,

AT2G17360, AT3G05590, AT4G02510, AT3G09500, AT2G21330, AT2G01140,

AT1G10510, AT4G33510, AT4G03280, AT5G09650, AT1G12410, AT2G23390,

AT3G47470, AT5G03470, AT3G12010, AT2G04390, AT1G09620, AT4G13940,

AT3G23810, AT3G08580, AT4G28390, AT1G73110, ATCG00350, AT5G14040,

AT3G48850, AT2G45960, AT1G06680, AT5G22800, AT5G42390, AT4G02930,

AT2G04520, AT1G71080, AT3G02830, AT3G04790, AT5G08050, AT2G37620,

AT1G02130, AT5G14720, AT5G38410, AT3G54890, AT3G26650, AT5G02160,

AT4G12800, AT3G55610, AT5G60790, AT3G54540, AT1G49240, AT2G42100,

AT1G78570, AT3G14415, AT5G17920, AT5G20980, AT1G04410, AT2G16570,

AT3G49910, AT1G43170, AT1G61580, AT2G25450, AT5G13780, AT2G27530,

AT1G67090, AT4G30010, AT1G20693, AT5G46800, AT2G03220, AT2G36580,

AT1G79850, AT1G56510, AT1G56520, AT2G16600, AT1G50200, AT1G29770,

AT5G42650, AT5G26742, AT1G09310, AT5G51970, AT1G33120, AT4G10450,

AT1G20260, AT1G09340, ATCG01060, AT1G63970, AT1G35720, AT1G06430,

AT1G03630, AT3G14310, AT1G53830, AT3G06510, AT4G29270, AT4G38630,

AT3G53110, AT1G70410, AT5G25780, AT1G20200, AT5G01410, AT3G10920,

AT1G74960, AT3G20820, AT3G02020, AT4G24190, AT2G13360, AT2G37270,

ATCG00340, AT3G57890, AT1G54340, AT3G15190, AT5G67030, AT3G47520,

AT5G19940, AT5G47435, AT1G34000, AT5G41210, AT2G01250, AT3G25520,

AT4G30190, AT1G17260, AT1G80660, ATCG00740, AT4G16720, AT3G17390,

AT4G34620, AT5G58140, AT1G62020, AT1G53750, AT3G08920, AT1G53500,

AT5G17170, AT5G27770, AT2G33800, AT2G20260, AT1G52360, AT1G23190,

AT1G70730, AT2G42740, AT1G48830, AT2G31610, AT5G35530, ATCG00520,

AT3G05560, AT2G44120, AT3G13580, AT2G40510, AT3G49010, AT5G15200,

AT5G39850, AT2G04842, AT1G27400, AT1G67430, AT3G10950, ATCG00840,

AT1G18500, AT3G04840, AT3G18740, AT1G70290, AT4G01310, AT3G07110,

AT4G13170, AT5G48760, AT2G21390, AT2G47610, AT3G52180, AT4G18100,

AT1G77940, AT3G53870, AT2G18450, AT4G31480, ATCG00770, ATCG00160,

AT1G66530, AT2G47730, AT2G02930, AT2G20450, AT1G09100, AT5G23540,

AT3G22110, AT2G38040, AT5G10360, AT3G45030, AT3G47370, AT2G03750,

AT2G19750, ATCG00820, AT1G09020, AT1G02780, AT4G02230, AT1G20950,

AT2G38140, AT3G57410, AT2G47940, AT3G13470, AT4G35090, AT1G20630,

ATCG00810, AT4G31700, AT2G33450, AT5G65010, AT5G10240, AT2G19740,

AT2G25670, AT5G28050, AT4G32470, AT3G08530, AT2G09990, AT3G04230,

AT4G30580, AT3G14120, AT3G56910, AT5G65220, AT1G63660, ATCG00065,

AT1G56110, ATCG00190, AT3G04870, AT5G28840, ATCG00500, AT1G19480,

AT1G30230, AT2G18110, ATCG00380, AT1G09200, AT1G13370, AT1G75600,

ATCG01120, AT1G09640, AT2G42790, AT2G41840, AT3G57490, AT2G05120,

AT4G31180, AT5G47190, AT4G17560, ATCG00900, AT5G26830, AT3G09630,

AT1G57720, AT1G06700, AT5G14320, AT1G10940, AT1G07920, AT3G57150,

AT3G46970, AT2G41530, ATCG00750, AT2G31810, AT1G72300, AT3G46520,

AT5G03100, AT5G12900, AT3G18780, AT3G13920, AT3G56340, AT5G56450,

AT5G60390, AT2G44060, AT5G42190, AT2G26790, AT1G02520, AT1G01320,

AT4G14960, AT4G03550, AT1G80870, AT1G70490, AT2G29420, AT3G58350,

AT5G19780, AT4G15200, AT1G09850, AT1G29900, AT1G02250, AT5G07090,

AT5G41140, AT2G45500, AT5G66080, AT1G58270, AT3G54560, AT3G11130,

AT3G51470, AT1G31180, AT1G64790, AT1G04560, AT5G07740, AT3G30842,

AT2G28890, AT3G10180, AT3G63420, AT5G42130, AT5G16760, AT5G65450,

AT3G25690, AT3G17650, AT1G79370, AT3G14660, AT1G03370, AT1G18440,

AT3G23175, AT4G37790, AT3G11940, AT5G23580, AT5G57390, AT1G73670,

AT5G44130, AT3G53510, AT3G20320, AT1G04640, AT3G16040, AT3G13410,

AT3G30820, AT4G29010, AT1G74070, AT3G24080, AT2G23200, AT5G13730,

AT3G28550, AT5G16740, AT5G46110, AT2G43480, AT2G24520, AT3G23450,

AT5G40240, AT5G19090, AT3G45980, AT1G47390, AT3G18900, AT3G47200,

AT2G21410, AT5G50000, AT5G59370, AT5G59700, AT2G15820, AT4G12420,

AT4G37900, AT5G63170, AT1G06990, AT5G46750, AT5G22490, AT1G67740,

AT2G46790, AT1G10010, AT3G62170, AT1G48220, AT1G01980, AT5G35670,

AT5G47020, AT5G49760, AT5G54350, AT1G11820, AT4G11730, AT5G01240,

AT1G61250, AT2G38290, AT5G67440, AT1G53210, AT1G36160, AT4G15530,

AT1G45160, AT1G75630, AT5G11110, AT1G05150, AT3G21820, AT5G57990,

AT2G35960, AT3G57810, AT4G13280, AT5G39190, AT2G39530, AT3G03310,

AT5G17320, AT2G39130, AT3G10440, AT4G08550, AT4G20130, AT2G31890,

AT4G16120, AT1G78200, AT3G53610, AT2G23250, AT3G54850, AT2G39480,

AT5G42880, AT4G26970, AT3G60240, AT1G74250, AT1G47900, AT2G11270,

AT5G02170, AT4G35230, AT1G15780, AT3G06740, AT3G25290, AT1G05880,

AT3G09260, AT4G31860, AT5G07910, AT1G68875, AT3G24300, AT4G23710,

AT3G52920, AT3G27700, AT3G61090, AT3G55410, AT5G51330, AT4G24630,

AT3G42640, AT4G34460, AT2G46110, AT3G27870, AT1G25350, AT1G52400,

AT4G09760, AT2G39330, AT3G19380, AT4G10490, AT5G01200, AT1G32050,

AT5G35735, AT4G02620, AT3G58730, AT3G46350, AT5G62740, AT3G61530,

AT3G08510, AT2G47000, AT1G69840, AT1G24490, AT3G16470, AT4G27585,

AT2G26730, AT1G13420, AT2G45820, AT3G01290, AT4G25960, AT3G28715,

AT3G61260, AT1G10290, AT3G53020, AT4G15470, AT3G55940, AT3G21090,

AT1G14320, AT4G11540, AT1G18310, AT5G42020, AT5G23400, AT5G06530,

AT4G01820, AT1G10680, AT4G29900, AT1G77210, AT1G70940, AT3G24550,

AT5G65700, AT1G28440, AT2G04080, AT5G42970, AT4G04700, AT2G37710,

AT3G17680, AT3G46270, AT5G04500, AT5G04430, AT1G26930, AT3G26520,

AT4G31710, AT3G48390, AT4G15620, AT1G13210, AT1G60030, AT1G58230,

AT1G52190, AT2G29110, AT1G23530, AT3G54140, AT4G38510, AT4G15610,

AT1G65200, AT2G41980, AT2G26975, AT1G12840, AT1G19110, AT5G06320,

AT2G22620, AT3G01460, AT3G54820, AT1G11260, AT3G28860, AT3G02880,

AT1G02530, AT4G18050, AT4G22120, AT3G14570, AT1G52191, AT3G07160,

AT5G61210, AT2G36850, AT3G09740, AT2G32450, AT5G27330, AT2G07702,

AT2G38460, AT2G07690, AT4G15630, AT4G28260, AT3G60330, AT5G16590,

AT3G08680, AT1G64200, AT1G59610, AT1G14830, AT5G43460, AT2G39250,

AT4G04370, AT3G01390, AT3G24160, AT1G73650, AT1G76030, AT1G79120,

AT1G05970, AT5G44790, AT2G36170, AT4G13510, AT3G17840, AT4G39080,

AT3G60190, AT3G06130, AT1G10050, AT3G01380, AT2G16850, AT1G30360,

AT1G48480, AT2G37180, AT4G12980, AT5G42080, AT3G61430, AT2G39010,

AT5G62670, AT2G18960, AT4G00430, AT1G01620, AT4G23400, AT4G35100,

## Acknowledgements

We thank Lisa Imrie and Katalin Kis for expert technical support. This work was supported by the Wellcome Trust [096995/Z/11/Z], by BBSRC and EPSRC awards [BB/N005147/1] [BB/D019621] and [BB/J009423], by NIH grants (GM079712 to T.I., NIH/NGMS P41 GM103533 to MJM) and by Next-Generation BioGreen 21 Program grant (SSAC, PJ011175, Rural Development Administration, Republic of Korea) to T.I. The authors declare no conflicting interests.

## Legends for supporting information

### Supplementary table

Table S1: Oligonucleotide primer sequences.

### Supplementary figures

**Figure S1**: Validation of the GI-TAP procedure in the preliminary study, showing the gel bands selected for mass spectrometric anlaysis.

**Figure S2**: Outlier analysis of the GI-TAP timeseries, showing statistical results that identified an outlier in sample 19E, which was removed from subsequent analysis.

**Figure S3**: Proteomics profiles of significantly enriched, plastid localized proteins.

**Figure S4**: The S-adenosyl methionine (SAM) cycle, showing 5-7 of the 9 genes listed that were enriched by GI-TAP.

### Supporting Experimental Procedures

**Methods S1.** Verification of GI:3F6H expression by Western Blotting and Silver-stain gel followed by in-gel digestion.

**Methods S2.** Protein extraction and enrichment by TAP for mass spectrometric analysis.

**Methods S3.** Protein digestion, peptide and mass spectrometric analysis.

**Overview of supporting data files [Note: .zip files that cannot be uploaded will be provided online upon publication]**

| Data file name | File contents |
| --- | --- |
| Data S1.xlsx | Proteins identified by LC-MS analysis of bands excised from silver-stained gel after GI-TAP (Preliminary study) |
| Data S2.xlsx | Proteins identified in the qualitative, in-solution digest analysis (Qualitative study) |
| Data S3.xlsx | Proteins identified in the timeseries study, resulting from the output of script R_stats_plots, with JTK_CYCLE results (Timeseries) |
| Data S4.xlsx | TopGO analysis results of GI-TAP timeseries |
| Data S5 |  |
| R_corr_PCA.R | R script for correlation analysis and PCA |
| corr_all.pdf | correlation analysis results for all GI samples, same as Fig. S2 |
| PCA_all.pdf | PCA on raw data, all samples |
| PCA_GI.pdf | PCA on raw data, only GI samples |
| PCA_GI_without19E | PCA on raw data, only GI samples and without outlier 19E, as Figure 2e |
| input_corr_PCA.csv | input file for correlation analysis and PCA |
| Data S6.zip |  |
| topGO_input_stats_raw.csv | stats input file for GO analysis (data taken from data S2 and JTK_CYCLE analysis) |
| topGO_output_allIDs.xlsx | direct output file of GO analysis for all identifications |
| topGO_output_min2IDs | direct output file of GO analysis for proteins with more than 1 identified peptides |
| gene-association.csv | GO database used for topGO analysis |
| Data S7.zip |  |
| R_stats_plots.R | R script for statistics and plotting |
| Progenesis_protein_measurements_raw.csv | Progenesis protein measurement output, raw abundance |
| Progenesis_protein_measurements_norm.csv | Progenesis protein measurement output, normalized abundance |
| R_stats_output | output file of script R_stats_plots.R |
| Data S8.zip |  |
| R_run_JTK_CYCLE.R | R script to run JTK_CYCLE analysis. The JTK_CYCLE resource must be in the same folder. |
| metadata.csv | metadata needed to run R_run_JTK_CYCLE |
| JTK_input.csv | input file into JTK_CYCLE analysis (Progenesis raw protein measurements output) |
| JTK_output.xlsx | output file from JTK_CYCLE analysis, was integrated into data S2 using Excel's vlookup function |
| Data S9.csv | Progenesis peptide measurements output file for timeseries study. |

## Supplementary Information

**Figure S1.**
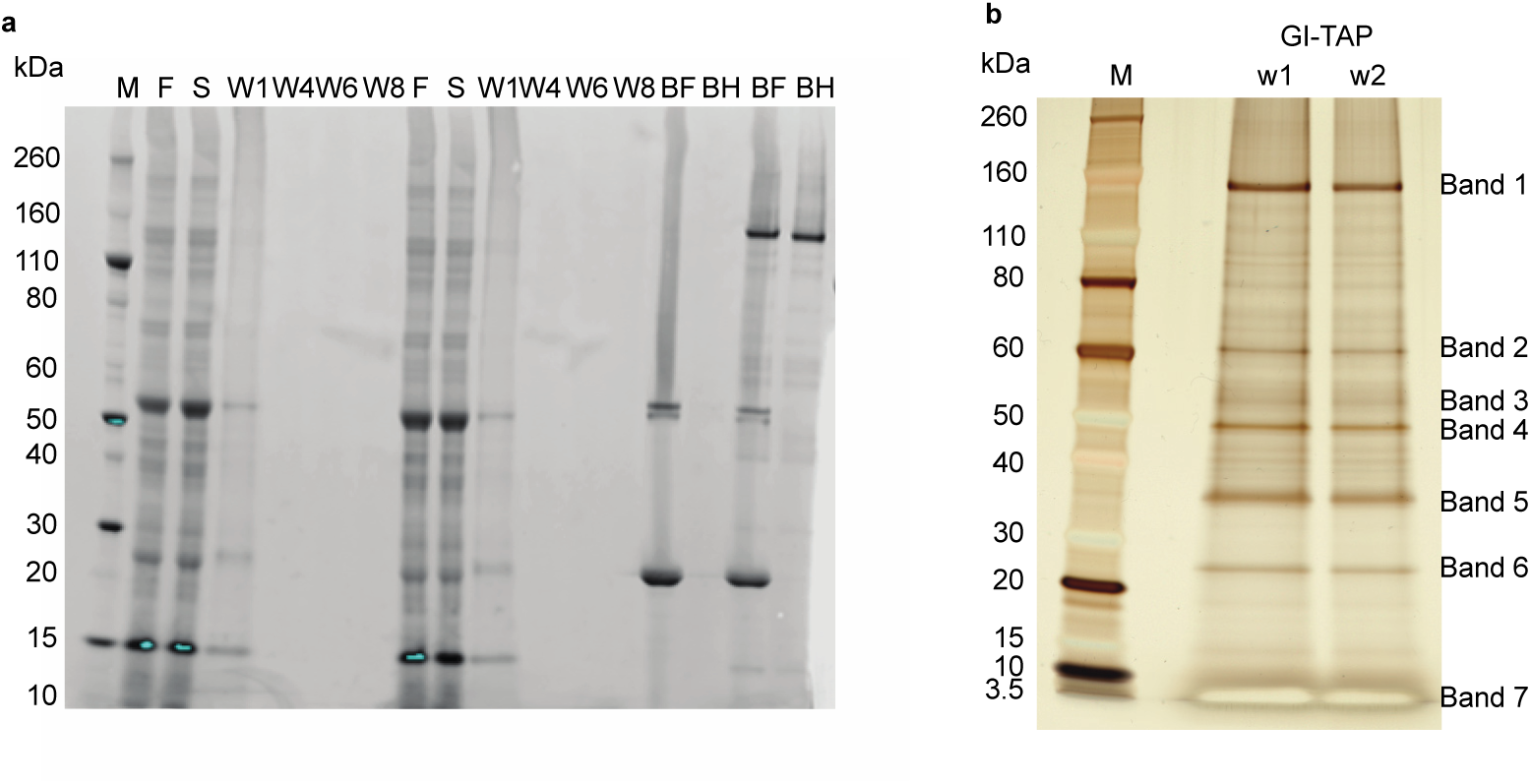
Validation of the GI-TAP procedure. (a) Protein samples from relevant steps of the GI-TAP procedure were analysed by immunoblotting, using an anti-Flag primary antibody and fluorescently-labelled secondary antibody, detected by fluorescence scanning. (b) Silver-stained gel of GI-TAP samples eluted from His-affinity matrix, showing bands tested by LC-MS (see Data S1; band 1 includes most GI; band 2 includes LKP2, FKF1 and ZTL, and so on). Gel track labels, M: Marker, F: filtered extract, S: supernatant after incubation with anti-Flag M2 beads, W1 to W8: washes, BF: Anti-Flag beads after incubation and before elution, BH: His-affinity matrix after incubation show purified GI:3F6H with a range of secondary bands. w1, w2: well number for MS analysis.

**Figure S2.**
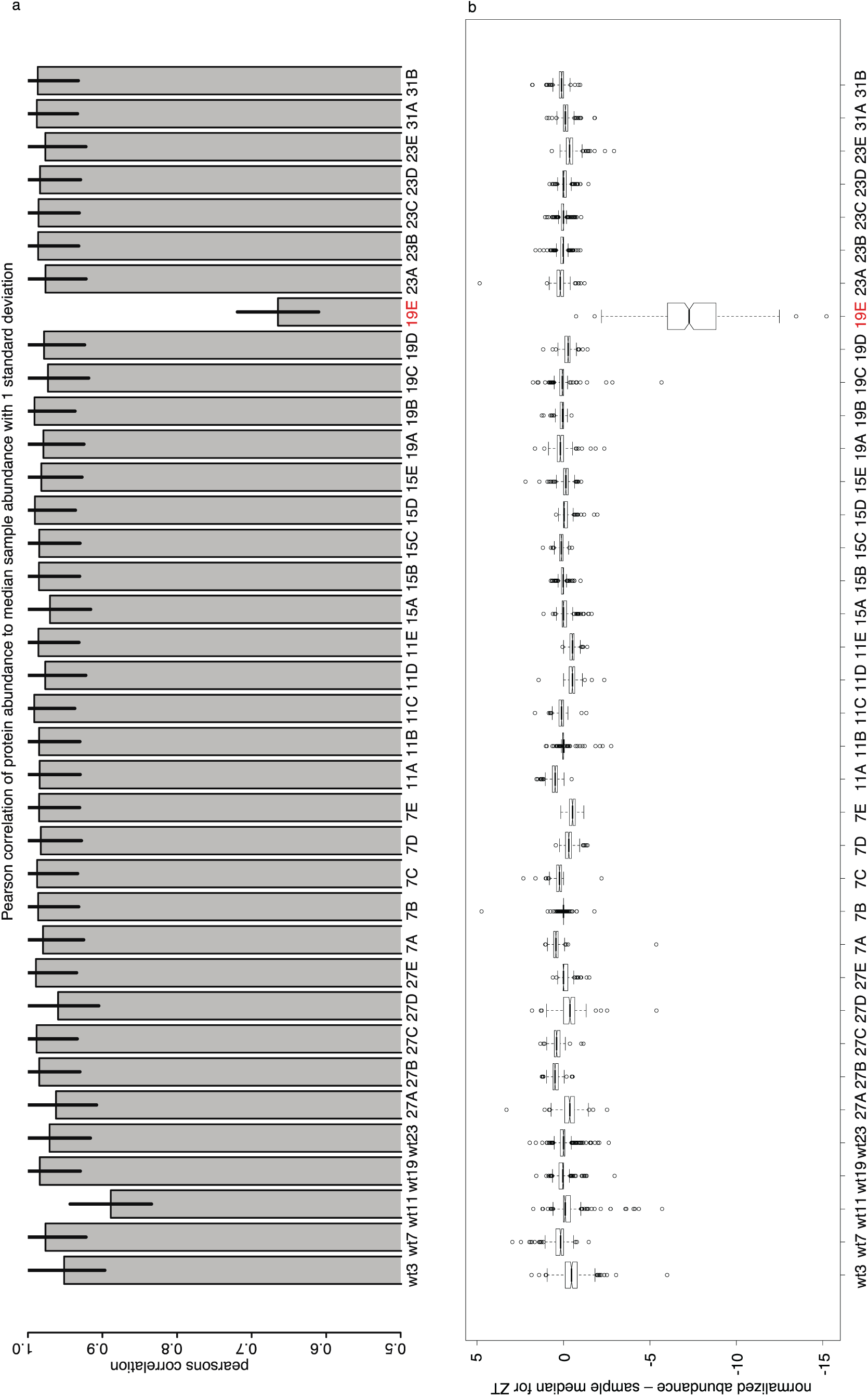
Outlier analysis of the GI-TAP timeseries study. Correlation analysis of the GI-TAP time course dataset, using analysis script in Data S5 to identify less-correlated outlier samples. a) Pearson’s correlation of protein abundance to median sample abundance, for each replicate. Error bars: standard deviation. b) Boxplot diagram of the differences between the protein abundance in individual replicate values and the median value at each timepoint, for each replicate. Outlier 19E is indicated in red.

**Figure S3.**
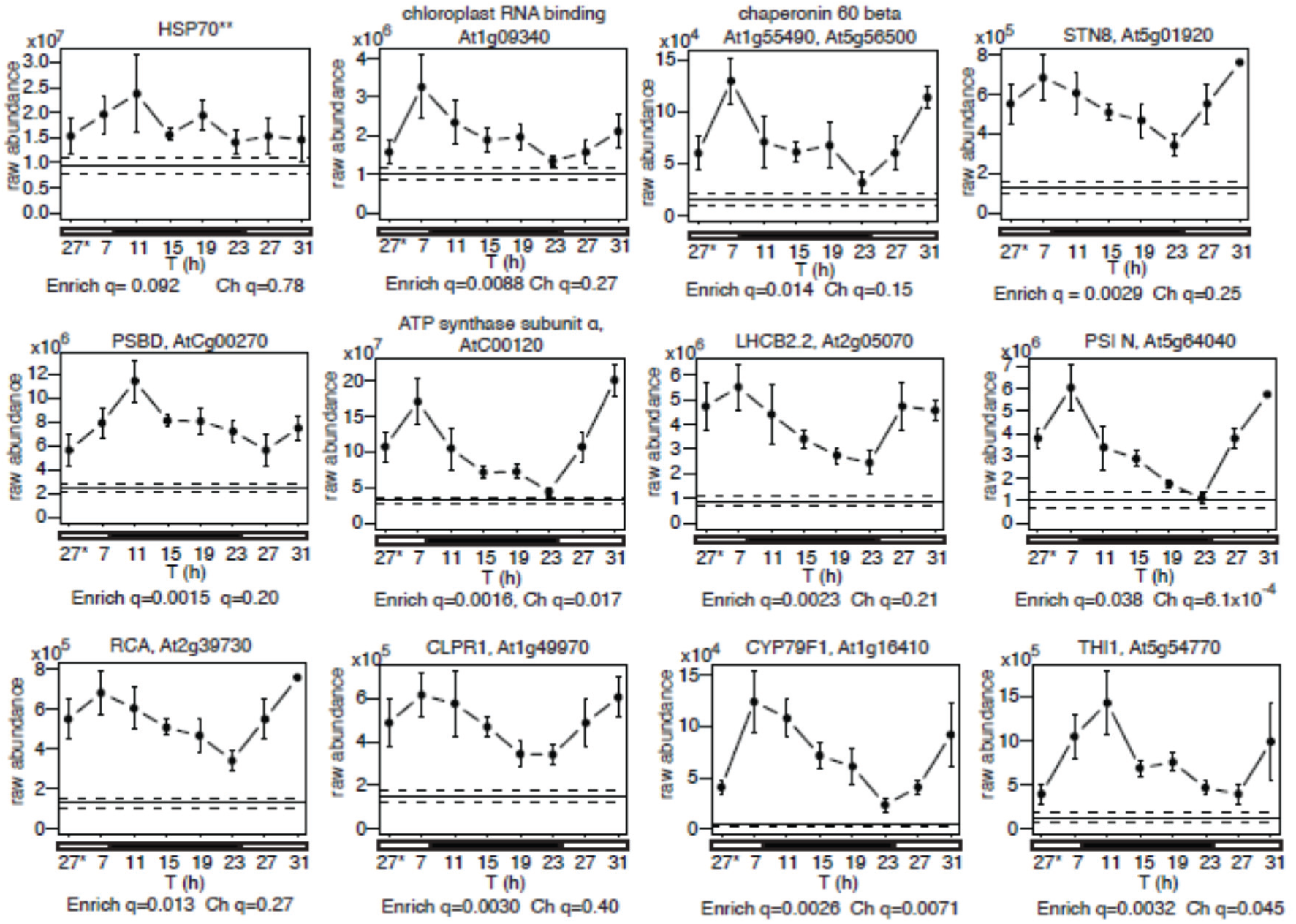
Diel profiles of HSP70 and significantly enriched, plastid localized proteins. Abundance profiles of HSP70 homologues (top left) and plastid-localized proteins. ** HSP70 proteins that were not distinguished by the eleven peptides detected: AT5G02500, AT1G16030, AT1G56410, AT3G09440, AT3G12580, AT5G02490, AT5G28540. GI-TAP samples, markers; error bar, SEM. Average of WT control, horizontal line, +/− SEM, dashed line. Time (T); * time point 27h is double-plotted. Significance of enrichment and temporal change are shown, as q-values of t-test comparing GI-TAP peak to WT (Enrich q) and of ANOVA within GI-TAP timeseries (Ch q).

**Figure S4.**
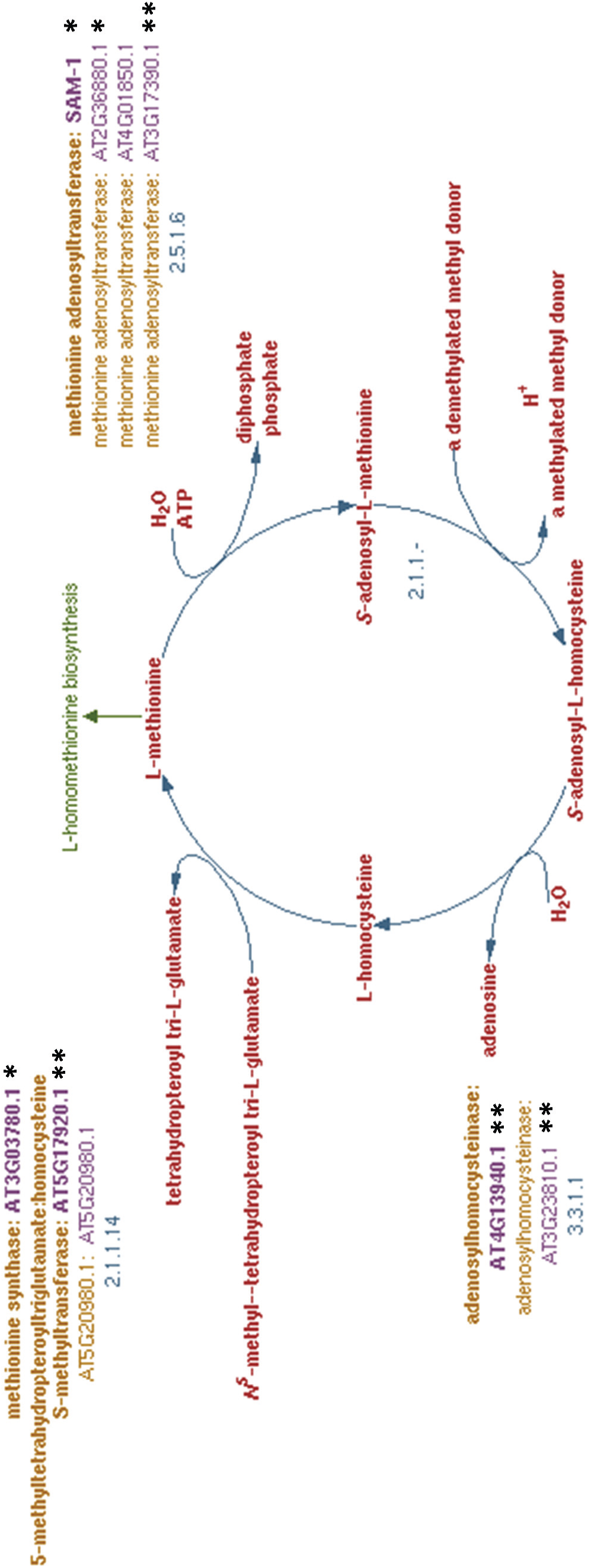
S-adenosyl methionine (SAM) cycle enzymes enriched by GI-TAP. Arabidopsis proteins annotated to the pathway (PlantCyc:PWY-5041) are highlighted by their identification in 1-2 GI-TAP interaction datasets (*-**), on the “S-adenosyl-L-methionine cycle II” pathway diagram from the PlantCyc resource (Plant Metabolic Network). 5-7 of the 9 genes listed were identified, as some identifications did not distinguish close homologues. The SAM-dependent methyltransferase (2.1.1-) and its target (methyl donor) are not specified.

## Supplementary tables

**Table S1.**
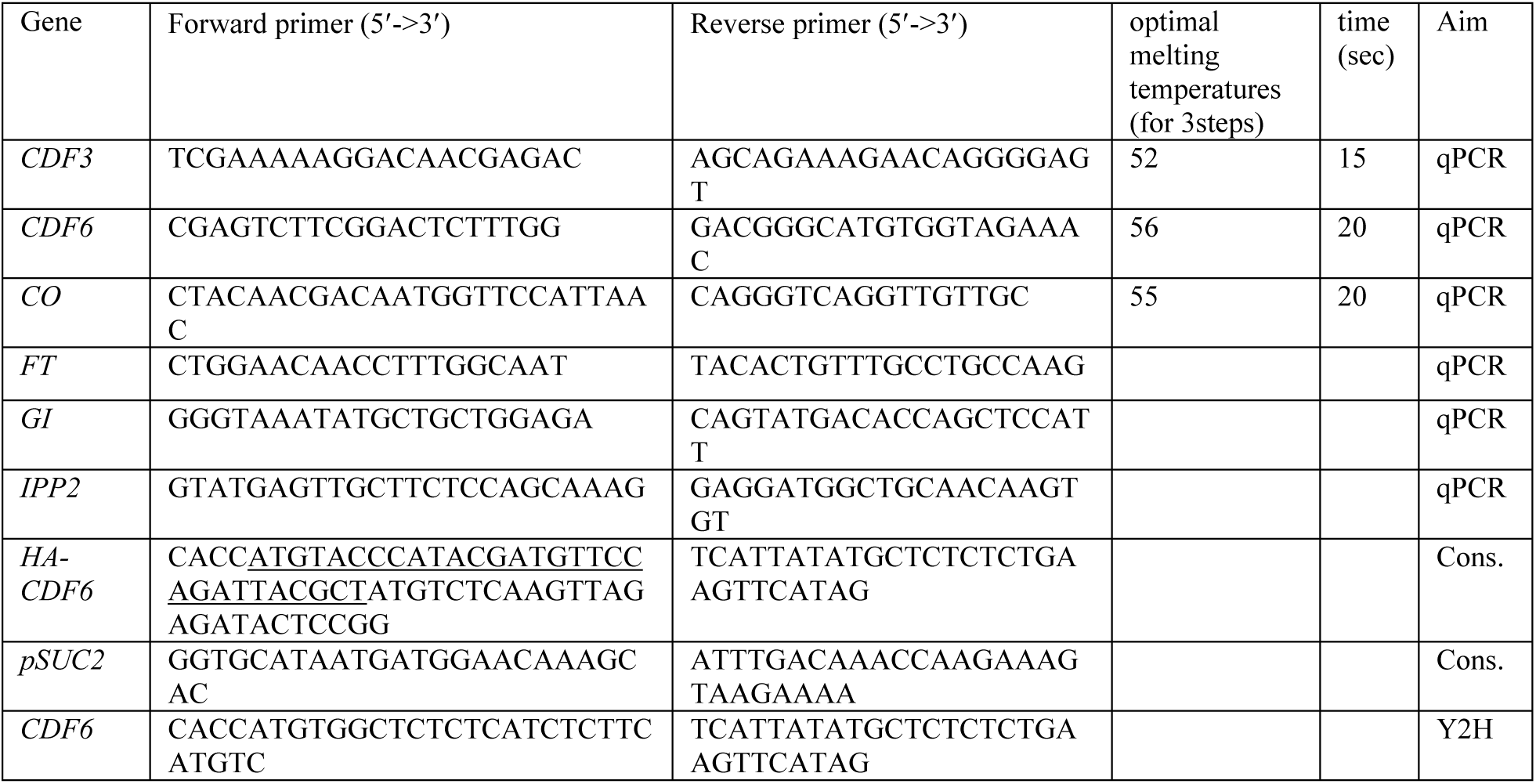
Primer sequences. Oligonucleotide primers were used for several purposes (Aim), qPCR RNA quantification, transgene construction (Cons.) and yeast two hybrid vector construction (Y2H). The HA epitope tag sequence fused to CDF6 is underlined.

## Supporting experimental procedures

### Methods S1

Verification of GI:3F6H expression by Western Blotting and Silver-stain gel followed by in-gel digestion

Western blot analysis shown in Figure S1 was performed as described (Song et al., 2014), using a commercial 4-12% Bis-Tris PAGE (Life Technologies) and the iBlot system according to the manufacturer’s instructions. The M2 mouse anti-Flag antibody (Sigma) and the IRDYE® 800 CW goat anti mouse secondary antibody (Licor) were used for detection of anti-Flag signal on the membrane using a Licor Odyssey fluorescence scanner. For silver staining, the SilverQuestTM silver staining kit (Life Technologies) was used according to the manufacturer’s instructions. Mass spectrometry results of in-gel digestion of specific bands are reported in data S1.

### Methods S2

Protein extraction and enrichment by TAP for mass spectrometric analysis Frozen plant tissue was ground to a fine powder in a liquid nitrogen and dry ice-cooled mortar and processed essentially as described (Song et al., 2014). For protein extraction, one tissue volume of SII buffer (100mM sodium phosphate pH7.4, 150mM KCl, 5mM EDTA, 5mM EGTA, 0.1% TritonX-100) or RIPA buffer (50mM Tris pH 7.5, 150mM KCl, 1% NP-40, 0.5% Dexoycholate) with 1 Complete protease inhibitor Cocktail EDTA free mini tablet (Roche) per 10ml, PhosStop phosphatase inhibitor mixture (Roche), 50µM MG-132 and 1mM PMSF was added to the tissue and tubes were vortexed vigorously. Crude extracts were sonicated with a sonicating probe at 10µM amplitude for 10s three times, cleared at 3220xg twice and filtered through 0.45µm syringe filters. Protein was quantified with a standard Bradford assay. The remaining TAP procedures and mass spectrometry were as described (Song et al., 2014).

For the timeseries, extract containing 28mg of protein was used for TAP. For AP on anti-FLAG M2 magnetic beads, a ratio of 10µl of beads (20µl of supplied 50% slurry, Sigma) per 4mg of protein was used. To equilibrate, beads were washed in ten packed gel volumes of TBS (50 mM Tris HCl, 150 mM NaCl, pH 7.4) twice, followed by one wash with SII or Ripa buffer without inhibitors. Beads were added to appropriate volume of extract as a 50% slurry and the mix was incubated on a rotating wheel at 4°C for 2h for binding of protein to the beads. Beads were washed twice with 20 bead volumes and once with ten bead volumes of SII buffer without inhibitors (1 bead volume being 100% bead gel), followed by two washes with 10 bead volumes of Flag-to-His buffer (0.5M NaPhosphate pH7.4, 150mM KCl, 0.05% TritonX-100). Protein was eluted 3 times with 2 bead volumes each of Flag-to-His buffer containing 250µg/ml 3xFLAG peptide (sigma) for 15min each. For 6xHis tag purification, 50µl of bead slurry were used for each sample. Dynabeads® for His purification (Life Technologies; timeseries) or Dyna1 Protein G magnetic beads (qualitative study) were equilibrated by washing with 20 slurry volumes of Flag to His buffer twice. Combined eluates were incubated with the beads on a rotating wheel for 15min. Beads were washed twice with 1ml of Flag-to-His buffer followed by three washes with 1ml of freshly prepared 25mM Ammonium Bicarbonate (Sigma). After removal of the final wash, beads were stored at −80°C until on-bead digest.

### Methods S3

Protein digestion, peptide and mass spectrometric analysis

For the qualitative study, the protein digestion, peptide analysis and mass spectrometry were as described (Song et al., 2014).

For the timeseries, on-bead digest was performed at room temperature as follows: 25µl 10mM DTT was added to the beads and samples were incubated at room temperature for 30min to reduce sulfhydryl groups. For carbamidomethylation, 50µl of 25mM iodoacetamide (sigma) were added and samples were incubated in the dark for 1h. 25µl of 8M urea, 2.5 µl of 1M ABC and 1.25µl of 1g/l trypsin (Worthington) were added and protein was digested overnight. An additional 0.25µl of 1g/l trypsin was added and digestion was performed for an additional 6h with occasional vortexing.

Peptides in the digest solution were separated from the Dynabeads® and desalted on a reverse phase resin using BondElute columns (25MG columns, Agilent): two times 1ml methanol and two times 1ml HPLC grade water (Thermo Fisher) were sequentially passed through the column before loading the sample. After binding of peptides, columns were washed using 1ml of HPLC grade water and peptides were eluted in 1ml acetonitrile. Peptides were vacuum dried in a Speed-vac (RC1010, Thermo). Dried peptides were dissolved in 8µl 0.05% TFA and passed through Millex-LH 0.45µm (Millipore) filters. 5µl were analysed by mass spectrometry. Nano-HPLC-MS/MS analysis was performed using an on-line system consisting of a nano-pump (Dionex Ultimate 3000, Thermo-Fisher, UK) coupled to a QExactive instrument (Thermo-Fisher, UK) with a pre-column of 300 µm x 5 mm (Acclaim Pepmap, 5 µm particle size) connected to a column of 75 µm x 50 cm (Acclaim Pepmap, 3µm particle size). Samples were analyzed on a 90min gradient in data dependent analysis (1 survey scan at 70k resolution followed by the top 5 MS/MS). One replicate of each time point was analyzed in random order before the next set of replicates of all time points, to rule out effects of instrument drift on quantitation.

## Appendix S1: Enrichment of Plastid proteins by GI-TAP

According to a Gene Ontology search using the Panther tool (data not shown), the timeseries study identified 166 plastid proteins; 101 of these were significantly enriched by at least two-fold. 10 or 11 were photosystem I or II components, respectively. Five proteins were annotated as light harvesting components. 53 proteins are localized in thylakoids, 35 in the stroma, nine in the chloroplast membrane. Three were plastid ATPase components. The more detailed, topGO analysis confirmed enrichment of plastid proteins by GI-TAP in the timeseries study (data S4). Among those plastid proteins were metabolic enzymes of the chloroplast: pyruvate kinase family protein (At3g22960), glyceraldehyde-3-phosphate dehydrogenase of plastid 2 (At1g16300) and fructose-bisphosphate aldolase 1 and/or 3 (AT2G21330, AT2G01140; data S3). In addition, some the photosynthetic proteins were enriched, such as LHCB2.2 (At2g05070), PSBD (AtCg00270), RCA (At2g39730) and PSI N (At5g64040), the protein kinase STN8 (At5g01920), the glucosinolate biosynthetic protein CYP79F1 (At1g16410) (Reintanz, 2001), and other functions such as plastid ATP synthase subunit (AtCg00120), and the thiamine biosynthetic gene THI1 (At5g54770). We also found enriched plastid proteins involved in protein folding and stabilization such as chaperonin 60ß, (At3g13470 and At1g55490 and/or At5g56500), CPN60A (At2g28000), a tetratricopeptide repeat family protein (At1g02150), HSP90.5 (At2g04030) and Hsp70 family protein TIC110 (At1g06950). The latter protein interacts with the plastid CLP protease family proteins, several of which were enriched, including CLPR1 (At1g49970), At1g09130 and CLPP5 At1g02560 (Figure S3, data S3).

